# MRCKα represses GEF-H1 mediated RhoA activation to promote ovarian cancer spheroid growth and invasion

**DOI:** 10.64898/2026.02.06.704384

**Authors:** Aasiya Remtulla, Alison M. Kurimchak, Alicia Canuel, Ishadi K. M. Kodikara, Jason S. Wasserman, Alexandre M. S. Chuyen, Danielle Olding, Rajesh Gupta, Nikolina Radulovich, James S. Duncan, Michael F. Olson

## Abstract

High-grade serous ovarian carcinoma (HGSOC) is characterized by high mortality rates and the frequent development of chemotherapy resistance. A hallmark of HGSOC progression is the formation of multicellular spheroids in malignant ascites that facilitate peritoneal dissemination and metastasis. While the CDC42-regulated kinases MRCKα and MRCKβ (MRCK) were previously found to be highly expressed in ovarian tumors and to be essential for cell migration and spheroid growth, the underlying molecular mechanisms were poorly defined. Mass spectrometry identified the RhoA-selective guanine nucleotide exchange factor GEF-H1 as a primary interacting partner of MRCKα. Functional assays revealed that MRCK inhibition or knockdown led to significantly increased levels of active GEF-H1 and subsequent RhoA activation. MRCKα was found to phosphorylate GEF-H1 on Ser174, and pharmacological inhibition of MRCK reduced this phosphorylation and decreased the association of GEF-H1 with α-Tubulin. Live-cell imaging and 3D assays demonstrated that MRCK inhibition disrupted cell-cell contacts and impaired the compaction of multicellular structures, ultimately reducing the viability of patient-derived organoids. These findings delineate a novel signaling crosstalk mechanism in which MRCKα represses GEF-H1-mediated RhoA activation to facilitate the formation of cell-cell contacts that contribute to the survival and growth of HGSOC cells in 3D multicellular structures. This study highlights MRCK as a potential therapeutic target to inhibit the growth and spread of HGSOC.

## Introduction

High-grade serous ovarian carcinoma (HGSOC) is one of the most common and lethal forms of ovarian carcinoma, and current treatments have only modestly impacted survival ^1^. The standard treatment for HGSOC is aggressive debulking surgery followed by platinum and taxane-based chemotherapy ^2^. Frequent alterations in DNA repair proteins such as BRCA1/2 sensitize HGSOC tumors to DNA-damaging agents, and to drugs that target homologous recombination ^1^. Although initial response rates are generally high, patients frequently develop chemotherapy resistance for which there is no current curative follow-up treatment ^3^. Moreover, the 5-year survival rate for HGSOC is abysmal at ∼40%, in part due to the high frequency of drug resistance and metastasis ^4^. Recurrent ovarian cancer cells often spread from primary tumors to the peritoneal cavity due to the lack of an anatomic barrier, resulting in disease dissemination ^5^. Malignant ascites, a buildup of fluid containing cancer cells in the abdomen, frequently occurs in late-stage HGSOC patients with recurrent disease ^6^. Tumor cells within the ascites of ovarian cancer patients often grow as spheroids or aggregates that are difficult to treat with conventional therapies and which enable the spread to other pleural organs, thus contributing to mortality ^5^. Consequently, new therapeutic strategies are urgently required for HGSOC, particularly therapies that inhibit tumor spheroid growth and survival, to reduce peritoneal and pleural metastasis.

Protein kinases (collectively termed the kinome) are arguably one of the most important types of regulatory enzymes in cells, and kinase inhibitory drugs have resulted in great improvements in cancer therapy ^7^. Despite their demonstrated importance and ready druggability, the contributions of many kinases to human diseases are poorly characterized, giving rise to the “dark kinome” concept that describes this lack of information for roughly one third of all kinases ^8^. A Multiplexed Inhibitor Beads and Mass Spectrometry (MIB-MS) approach was used to map the kinome of HGSOC primary and patient-derived xenograft (PDX) tumors, which led to the identification of the <u>M</u>yotonic dystrophy-<u>R</u>elated <u>C</u>dc42-binding <u>K</u>inases MRCKα and MRCKβ (collectively, MRCK) as being highly expressed in ovarian tumors ^9^. MRCK knockdown or pharmacological inhibition with the potent and selective inhibitor BDP9066 ^10^ demonstrated that MRCK signalling was essential for ovarian cancer cell migration, and the growth of established cell lines in 2D monolayers or as 3D spheroids^9^.

MRCKα and MRCKβ are members of the AGC kinase subfamily ^11,12^, and are CDC42 GTPase effectors that phosphorylate several substrates that contribute to the dynamic regulation of actin cytoskeleton organization including regulatory myosin light chains (MLC) ^13^, the myosin-binding subunit of the MLC phosphatase complex (MYPT1) ^14^, LIM kinase 1 (LIMK1) ^15^ and myosin 18A (MYO18A) ^16^. Elevated expression of the gene encoding MRCKα (*CDC42BPA*) was previously found to be associated with a short interval to distant metastasis in breast cancer patients ^17^. Consequently *CDC42BPA* has been included in the MammaPrint 70 gene set that is used as a diagnostic tool to predict the risk of breast cancer metastasis ^18^. Furthermore, in mouse pre-clinical cancer models the MRCK inhibitor BDP9066 significantly decreased chemically-induced skin cancer ^10^ and triple negative breast cancer xenograft growth ^19^, and reduced radiotherapy-induced spread of glioblastoma cells to significantly improve the survival of tumor-bearing mice ^20^. Despite the strong evidence linking MRCK with cancer, relatively little is known about the molecular mechanisms that contribute to tumor growth and progression. In fact, the considerable overlap in substrates of MRCK^11,12^ and RhoA-regulated ROCK kinases ^21^ obscures the unique aspects of MRCK function that promote cancer.

We now report that the selective MRCK inhibitor BDP9066 potently inhibited the proliferation of HGSOC patient-derived cells grown as three dimensional (3D) spheroids or organoids. Mass spectrometry identified the RhoA-selective guanine nucleotide exchange factor GEF-H1 as the most abundant MRCKα-interacting protein, and MRCK knockdown/knockout or pharmacological inhibition increased the pool of active GEF-H1 available for binding to nucleotide-free RhoA. Furthermore, MRCK knockdown/knockout or pharmacological inhibition led to RhoA activation in a GEF-H1-dependent manner. MRCKα phosphorylated GEF-H1 on Ser174, and pharmacological inhibition reduced the extent of S174 phosphorylation on active GEF-H1, which was paralleled by decreased association of GEF-H1 with α-Tubulin. Live cell imaging revealed that MRCK inhibition disrupted cell-cell contacts and promoted paired cells to move apart, effects that were blocked by GEF-H1 knockdown. Taken together, these observations delineate a novel mechanism that mediates cross-talk between Rho GTPase signaling pathways, which enables the formation of cell-cell contacts that facilitate the growth of HGSOC cells in multicellular 3D structures.

## Results

### Therapeutic effects of MRCK inhibition on ovarian cancer

Consistent with the observed elevation in MRCKα expression in ovarian cancer tumors ^9^, the overall survival of stage 4 ovarian cancer patients was significantly (logrank *p* = 0.0082) reduced from 35 months for patients with less than median expression to 25.3 months for patients with higher than median expression of *CDC42BPA* determined from publicly available Affymetrix gene expression databases (**Figure 1A**) ^22^. Tumor cells within the ascites of ovarian cancer patients often grow as spheroids, which are difficult to treat and frequently metastasize to other pleural organs ^6^. Spheroids form via sequential aggregation/association and compaction steps ^23^. Actin cytoskeleton organization is regulated by Rho GTPases, with the canonical Rho GTPases Rac1 and CDC42 contributing to the generation of membrane protrusions that enable the formation of cell-cell contacts ^24^, and RhoA promoting actin-myosin contractile force ^25^. Importantly, RhoA activity must be down-regulated at cell-cell contacts to permit the formation of 3D cell structures ^26^, consistent with excessive RhoA activity leading to the destabilization of epithelial cell-cell contacts ^27^. Latterly, RhoA activity ^28^ and actin-myosin contraction are necessary for reinforcement of cell contacts and spheroid compaction ^23,29–31^. Thus, spheroid formation and growth require precisely coordinated regulation of Rho GTPases. In prior work, we showed that treatment of established HGSOC cell lines with a pharmacological MRCK inhibitor impaired actin remodelling and reduced spheroid formation ^9^. Here, using a cell line derived from the ascites fluid of a HGSOC patient with recurrent disease (OC-1), we explored the impact of MRCK inhibition on the phosphorylation of proteins that influence actin-myosin remodelling, and cell viability and invasion in three-dimensional (3D) contexts. Consistent with previous findings ^10^, treatment of OC-1 cells with 5 µM BDP9066 for 4 hours significantly (*p* = 0.009) inhibited phosphorylation of MLC on Thr18/Ser19 (pMLC), and MYPT1 on Thr696 (*p* < 0.0001, pMYPT1) (**Figure 1B**). In 3D non-adherent growth assays, treatment of OC-1 cells with BDP9066 inhibited cell viability, exhibiting a GI_50_ = 6.89 µM (**Figure 1C**). The metastasis of ovarian cancer spheroids in the peritoneal cavity relies on the ability of the tumor cells to invade the mesothelium that covers peritoneal and pleural organs ^32^, a process shown to be dependent on actin-myosin generated force ^33^. To model this process, the ability of OC-1 spheroids to invade collagen was assayed. After 24 hours, OC-1 spheroids embedded in collagen that were treated with DMSO vehicle control had significantly (*p* < 0.0001) invaded and spread (**Figures 1D**). In contrast, treatment with 10 µM BDP9066 blocked collagen invasion such that it was not significantly different (*p* = 0.4133) from starting spheroid areas, and which was significantly (*p* < 0.0001) reduced relative to DMSO treated control cells (**Figures 1D**). Using a “reverse” collective cell invasion assay ^34^ in which OC-1 cells migrated through a porous filter and invaded into three-dimensional Matrigel, confocal microscopy revealed that treatment with 3 μM BDP9066 for 5 days reduced the invasion of mCherry-CAAX-expressing OC-1 cells into sequential 10 μm optical slices (**Figure 1E**). By quantifying the number of mCherry-CAAX fluorescent positive pixels at each 10 μm interval in three replicate experiments (**Figure 1F**), it became apparent that 3 μM BDP9066 significantly (*p* = 0.0058) reduced cell density (**Figure 1G**) and the proportion of cells that invaded ≥ 40 μm (*p* = 0.0262) (**Figure 1H**) into the 3D Matrigel. Thus, MRCK inhibition affected both the proliferation/survival and invasion of patient-derived HGSOC cells in 3D contexts.

**Figure 1.**
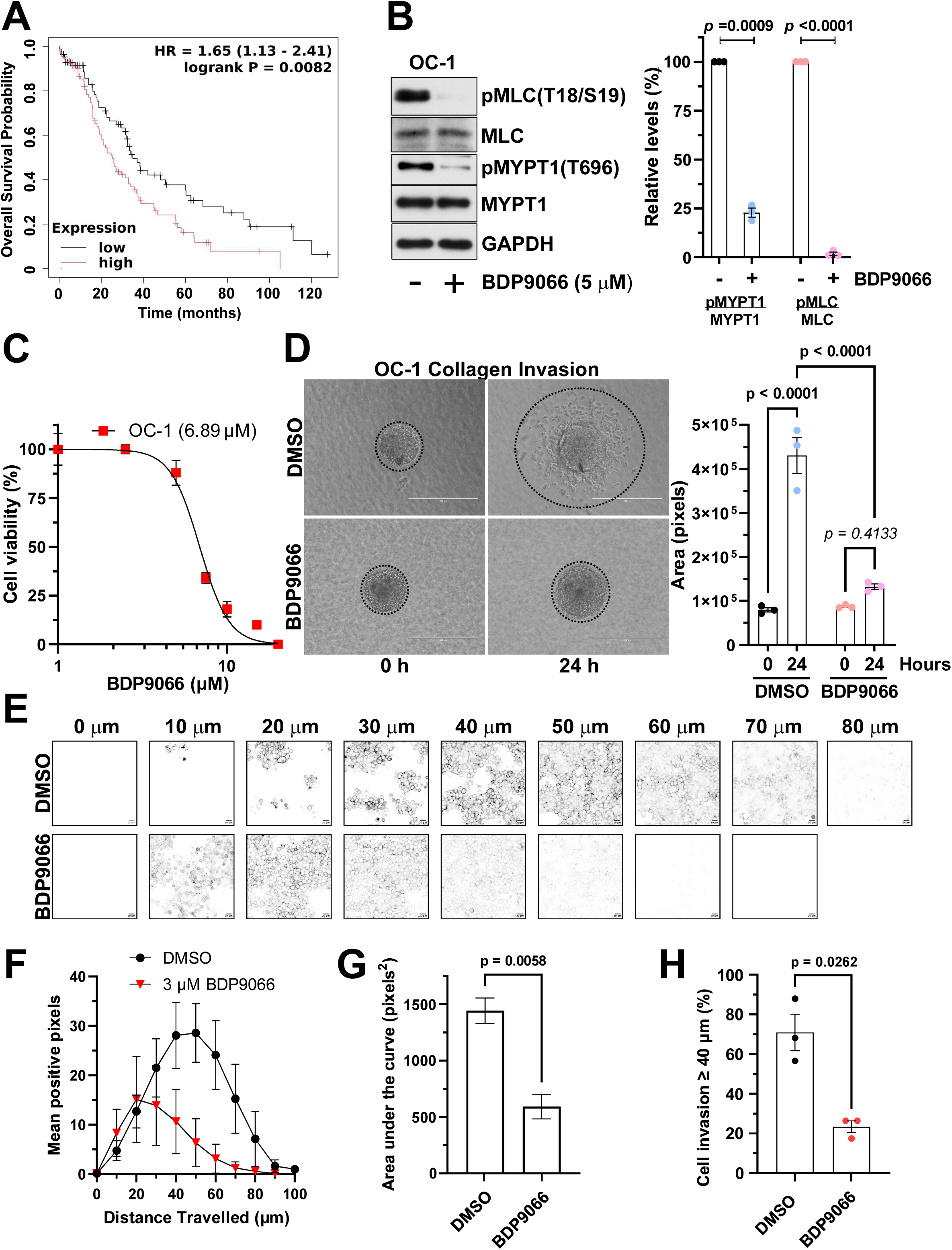
MRCK inhibition decreases patient-derived spheroid viability and invasion. **A**. Kaplan-Meier of the probability of overall survival over time for 176 stage 4 ovarian cancer patients determined from Affymetrix microarray gene expression data using the Jetset ^75^ prioritized probe 203794_at for *CDC42BPA.* Statistical analysis used logrank test to produce the displayed *p* value. **B.** (*Left panel*) Representative immunoblots of OC-1 cells treated with DMSO or 5 µM BDP9066 for 4 hours. (*Right panel*) Densitometric analysis of bands are means ± SEM from three independent replicates, each normalized to the GAPDH loading control followed by normalization of the phosphorylation relative to non-phosphorylated bands. Statistical analysis used unpaired t-tests with Welch’s correction producing the displayed *p* values. **C**. CellTiter-Glo assays of the viability OC-1 cells grown as 3D spheroids treated with increasing concentrations of DMSO or BDP9066. Data were analysed as percentages of DMSO control, presented as means ± SEM of three independent replicate assays. GI_50_ value was determined using log(drug) versus response using four-parameter variable slope nonlinear curve fitting. **D.** (*Left panel*) Representative images of OC-1 spheroids grown in type I Collagen were treated with DMSO or 10 µM BDP9066 for 24 hours. Scale bars = 400 µm. (*Right panel)* Spheroid invasion into collagen was quantitated in ImageJ by measuring the total area of invasion surrounding the spheroid. Area analysis of invasion are means ± SD from three images. Statistical analysis used unpaired t-test with Welch’s correction producing the displayed *p* values. **E.** Inverted fluorescence confocal microscope images taken at sequential 10 μm optical slices of mCherry-CAAX expressing OC-1 cell invasion into three dimensional Matrigel. Cells were treated with DMSO vehicle control or 3 μM BDP9066 for 5 days. Scale bars = 20 μm. **F.** Quantification of the number of mCherry-CAAX OC-1 cell positive pixels in 512 X 512 pixel images at each 10 μm optical slice up to 100 μm. Data are presented as means ± SEM of three independent replicate assays. **G.** Mean areas under the curve (AUC) were determined for DMSO and 3 μM BDP9066 treated OC-1 cells. AUC results are means ± SEM from three replicate assays. Statistical analysis used unpaired t-test with Welch’s correction producing the displayed *p* values. **H.** Percentage of cell invasion ≥ 40 μm of DMSO or 3 μM BDP9066 treated OC-1 cells. Results are means ± SEM from three replicate assays. Statistical analysis used unpaired t-test with Welch’s correction producing the displayed *p* values.

### MRCKα interacts with and regulates the RhoA selective GEF-H1 GDP/GTP exchange factor

To date, a detailed characterization of MRCKα interacting proteins has not been performed in any cell type. Notably, the recent large-scale analysis of the Ser/Thr kinome interactome did not analyse MRCKα ^35^. To identify MRCKα interacting proteins in HGSOC cells, we expressed FLAG-tagged MRCKα or GFP (control) in OC-1 cells and performed FLAG immunoprecipitations followed by Liquid Chromatography-Tandem Mass Spectrometry (LC-MS/MS). More than 1000 proteins were enriched in FLAG-MRCKα immunoprecipitation from OC-1 cells (**Figure S1A, Data File S1**), which many significantly differing from those enriched in FLAG-GFP immunoprecipitations (**Figures S1B-C**). Heat map (**Figure 2A**) or volcano plot analysis (**Figure S1D**) identified proteins that were uniquely enriched with FLAG-MRCKα relative to the control FLAG-GFP in OC-1 cells. Pathway analysis of proteins enriched in FLAG-MRCKα immunoprecipitations using Metascape revealed that these proteins were predominantly associated with Signalling by Rho GTPases, PTEN Regulation and Actin Cytoskeleton Organization (**Figure 2B**). Protein-protein interaction network analysis of proteins enriched Rho GTPase signalling using STRING ^36^, revealed several novel MRCKα interacting proteins, as well as established MRCKα interacting partners including CDC42 (**Figures 2C and S1E**) ^11^. Interestingly, the Rho Guanine Exchange Factor GEF-H1 (ARHGEF2) was amongst the top 5 most abundant proteins pulled down with FLAG-MRCKα based on spectral counts or Label-Free Quantification (LFQ) intensity (**Figures 2C and S1F-G**). GEF-H1 activates RhoA by promoting the exchange of GDP for GTP, and has been shown to be essential for proliferation, migration, and invasion in several cancer models ^37^. The FLAG-MRCKα interaction with GEF-H1 was confirmed by western blotting, in which endogenous GEF-H1 co-immunoprecipitated with FLAG-MRCKα or endogenous MRCKα co-immunoprecipitated with FLAG-GEF-H1 in OC1 cells grown as 2D monolayers (**Figure 2D**). Furthermore, the interaction of FLAG-MRCKα with endogenous GEF-H1 was validated when OC-1 cells were grown as 3D spheroids (**Figure 2E**).

**Figure 2.**
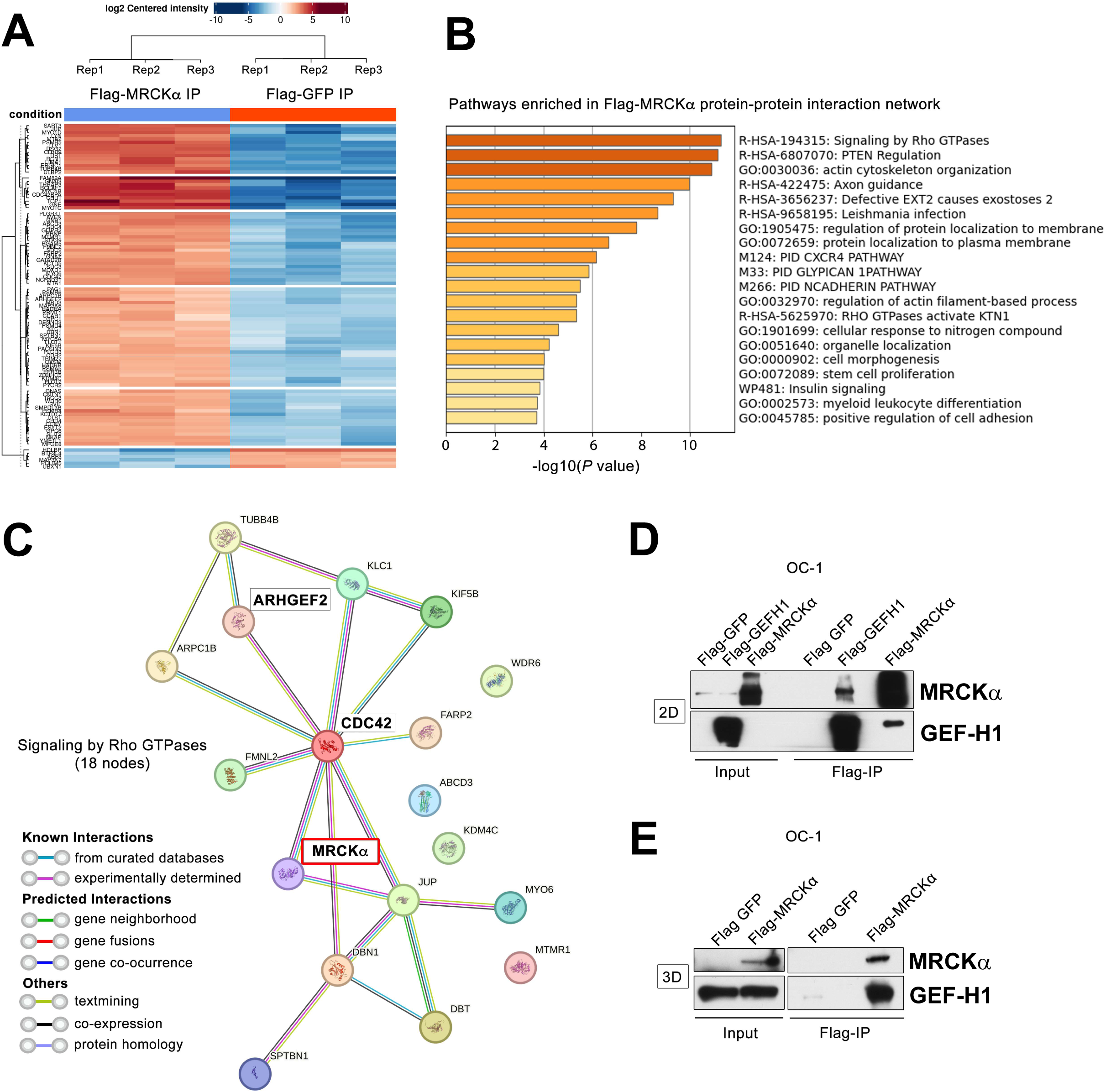
Characterization of MRCKα interacting proteins in HGSOC cells reveals a novel interaction with GEF-H1. **A**. Heat map depicts all significant proteins exhibiting differential abundance (rows) in FLAG-MRCKα immunoprecipitations relative to FLAG-GFP (columns) in OC-1 cells. OC-1 cells expressing FLAG-MRCKα or FLAG-GFP were subjected to FLAG immunoprecipitation in three independent biological triplicates and protein interactions were identified by mass spectrometry. Differences in log_2_ LFQ intensities among FLAG-MRCKα or FLAG-GFP immunoprecipitations were determined by paired t tests with an adjusted *p* value cutoff of 0.001 and log_2_ fold change cutoff of 4 using Benjamini Hochberg FDR correction. Analysis was performed using Phospho-Analyst ^76^. **B**. Bar graph depicts pathway enrichment analysis of proteins pulled down with FLAG-MRCKα but not FLAG-GFP immunoprecipitation in OC-1 cells. Proteins significantly enriched in FLAG-MRCKα immunoprecipitations (FDR < 0.05, *p* < 0.001, log_2_ fold cutoff = 4) were imported into Metascape pathway analysis. Bar graph colour depicts log_10_ *p* values for pathway enrichment. **C**. FLAG-MRCKα protein-protein interaction network enriched in signalling by Rho GTPases. Network image generated using String Version 12.0. Immunoprecipitation (IP) of exogenous FLAG-GFP, FLAG-MRCKα or FLAG-GEF-H1 from OC-1 cells grown in **D.** 2D or **E.** 3D as spheroids. Input samples and FLAG IPs were blotted with MRCKα or GEF-H1 antibodies as indicated.

To further validate the MRCKα interaction with GEF-H1 in ovarian cancer cells, reciprocal immunoprecipitations were performed with antibodies against MRCKα, GEF-H1 or MRCKβ, or a non-specific rabbit IgG, using A2780 endometroid adenocarcinoma, SK-OV-3 ovarian serous cystadenocarcinoma, OVCAR8 or OC-1 HGSOC cell lysates (**Figures 3A and S2A**). While immunoprecipitated endogenous MRCKα co-precipitated with GEF-H1 and *vice versa*, there was no evident MRCKβ co-precipitated with immunoprecipitated GEF-H1, nor did immunoprecipitated MRCKβ co-precipitate with GEF-H1, consistent with the GEF-H1 interaction being selective for MRCKα. Similar co-immunoprecipitation of endogenous MRCKα and GEF-H1 was observed for HEK293 human embryonic kidney and MDA-MB-231 human breast cancer cells (**Figure S2B**).

**Figure 3.**
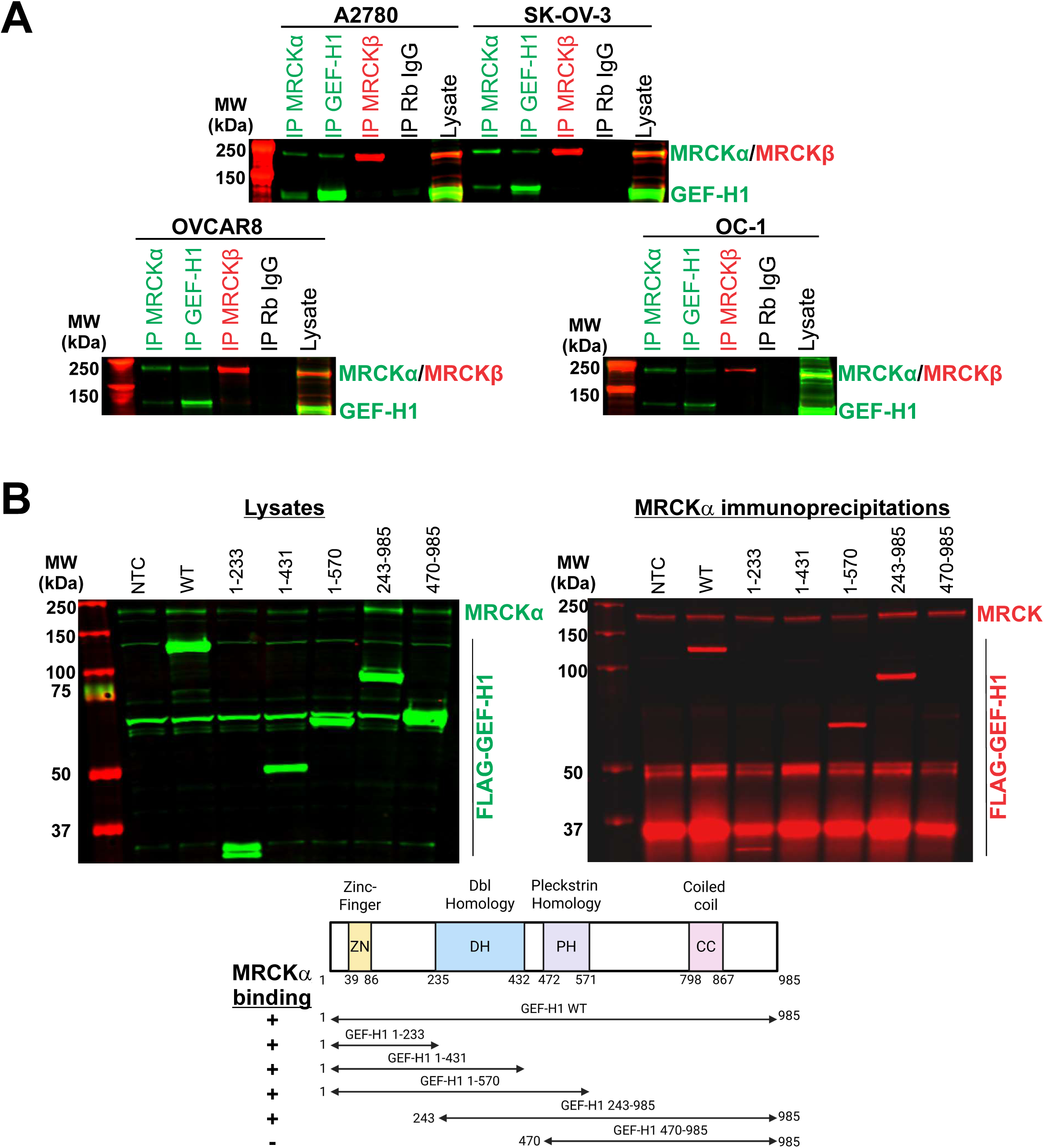
MRCKα interacts with the GEF-H1 N-terminus. **A.** Lysates from A-2780, SK-OV-3, OVCAR8 and OC-1 cells were incubated with antibodies against MRCKα, GEF-H1 or MRCKβ, or with non-specific rabbit IgG as indicated. Immunoprecipitated (IP) proteins and cell lysates were western blotted with antibodies again MRCKα (green), MRCKβ (red) and GEF-H1 (green). **B.** OC-1 cells were either non-transfected (NTC) or transfected with FLAG-tagged wild-type (WT) or deletion mutants that comprised the indicated amino acids. *Left panel:* Cell lysates were western blotted with antibodies against MRCKα (green) and FLAG-epitope tagged GEF-H1 (green). *Right panel:* Cell lysates were incubated with antibodies against MRCKα, and immunoprecipitated proteins were western blotted with antibodies against MRCK (green) and FLAG-epitope tagged GEF-H1 (green). *Lower panel:* Schematic diagram of GEF-H1 domains with a summary of GEF-H1 binding to MRCKα deletions. Zn – Zinc finger. DH – Dbl Homology. PH – Pleckstrin Homology. CC- Coiled Coil.

OC-1 HGSOC cells were transfected with FLAG-epitope tagged wild-type (WT) GEF-H1 or a series of deletion mutants ^38^ to map regions interacting with MRCKα. Each exogenous GEF-H1 protein was expressed at equivalent levels, and immunoprecipitation of endogenous MRCKα revealed binding of full-length WT MRCKα and mutants containing amino acids 1-243, 1-431, 1-570, 243-985 but not 470-985 (**Figures 3B and S2C**). These results are consistent with there being two separate regions, 1-243 and 243-470, that interact with MRCKα.

To investigate a possible link between MRCK and GEF-H1 regulation, OVCAR8 cells were transduced with lentivirus encoding the Cas9 nuclease and non-targeting sgRNA (NTC), or a combination of lentivirus encoding Cas9 and sgRNAs targeting MRCKα and MRCKβ to knockout expression (αβ KO). Quantitative western blotting determined that the knockout efficiency of both kinases in transduced cell pools was > 75% (**Figure S3A**). Activated RhoGEFs have higher affinity for nucleotide-free RhoA than for GDP-RhoA or GTP-RhoA ^39^. Previous studies used glutathione-S-transferase (GST) fused to RhoA with a G17A point mutation that blocks nucleotide binding to affinity purify active GEF-H1 from T47D human breast cancer and normal murine mammary gland cells that had been plated on matrices of varying stiffnesses ^40^. When GEF-H1 was affinity-purified with GST-RhoA G17A from NTC or αβ KO OVCAR8 cells, there was significantly (*p* = 0.0382) 37% more active GEF-H1 available for binding to nucleotide free RhoA in the αβ KO cells relative to NTC controls (**Figures 4A and S3B**). Pharmacological MRCK inhibition of OVCAR8 cells with 1 µM BDP9066 for 1 h similarly produced a significant (*p* = 0.0023) 75% increase in active GEF-H1 binding to RhoA G17A relative to DMSO vehicle treated control cells (**Figures 4B and S3C**). Similarly, 1 µM BDP9066 treatment resulting in significantly greater GEF-H1 being associated with GST-RhoA G17A in lysates from A2780 cells (**Figure 4C**, *p* = 0.0184, 110% increase) and OC-1 cells (**Figure 4D**, *p* = 0.0319, 38% increase). Taken together, these observations demonstrate that MRCKα interacts with GEF-H1, which influences its activation state and consequent ability to interact with nucleotide-free RhoA.

**Figure 4.**
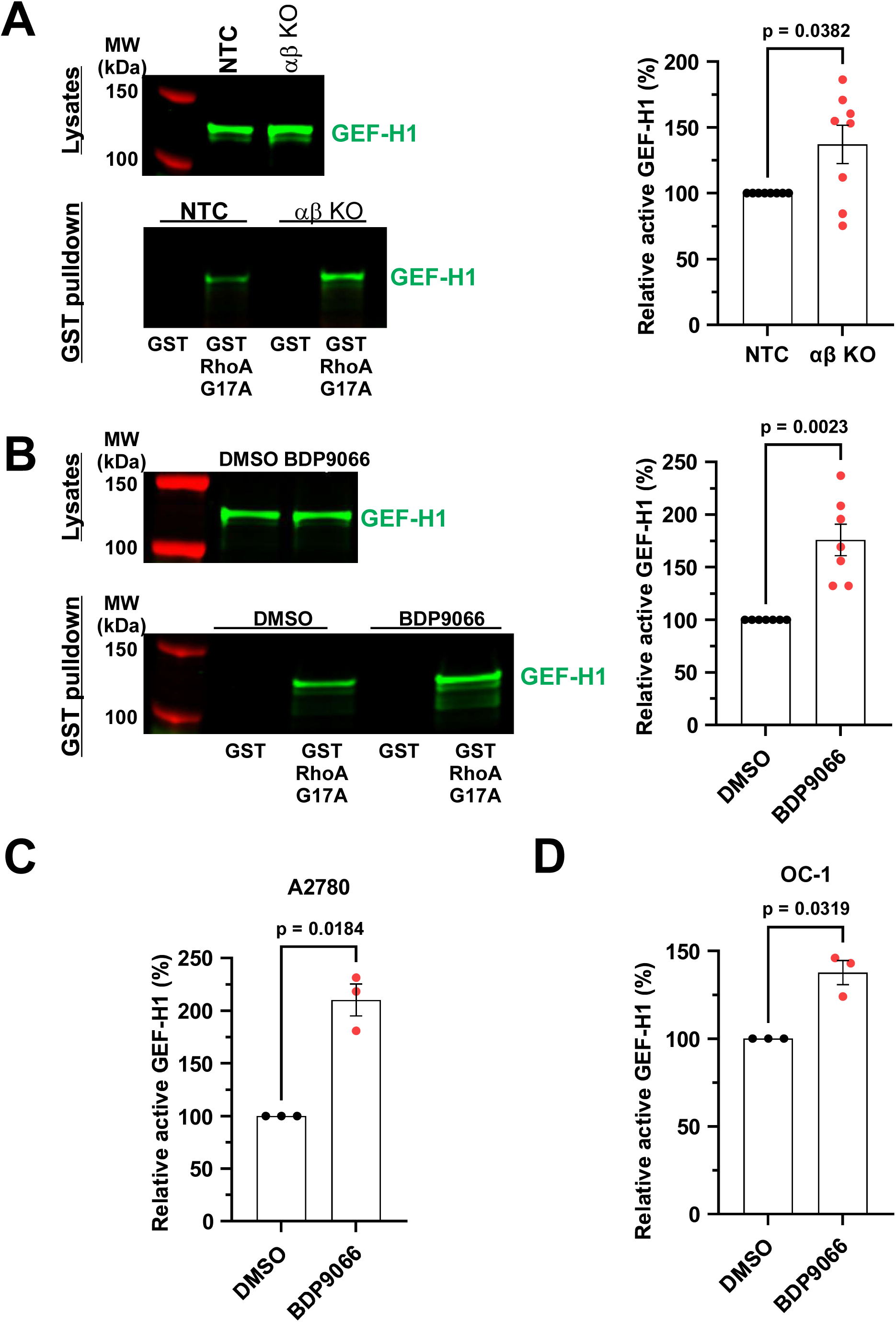
MRCK deletion or inhibition increases active GEF-H1 available for binding to nucleotide-free RhoA. **A.** *Left panel:* NTC or αβ KO OVCAR8 cells were lysed and western blotted with antibody against GEF-H1 (green). OVCAR8 NTC or αβ KO lysates were incubated with purified GST or nucleotide free mutant GST-RhoA G17A protein immobilized on reduced glutathione-conjugated Sepharose beads as indicated to pull-down active GEF-H1 available for binding to nucleotide-free RhoA. Washed glutathione Sepharose beads with immobilized GST fusion proteins associated with GEF-H1 were quantitatively western blotted with antibodies against GEF-H1 (green). *Right panel:* The quantified signal for GST-RhoA G17A associated GEF-H1 from αβ KO cells was reduced by the non-specific GEF-H1 signal associated with GST and the difference was divided by the input GEF-H1 signal, and the resulting ratio was normalized to the equivalent ratio from NTC cells for each independent replicate. Means ± SEM, N = 8. **B.** *Left panel:* OVCAR8 cells were treated with DMSO vehicle control or 1 µM BDP9066 for 1 h, as indicated, and were lysed and western blotted with antibody against GEF-H1 (green). DMSO or BDP9066 treated lysates were incubated with purified GST or nucleotide free mutant GST-RhoA G17A protein immobilized on reduced glutathione-conjugated Sepharose beads as indicated. Washed glutathione Sepharose beads with immobilized GST fusion proteins associated with GEF-H1 were quantitatively western blotted with antibodies against GEF-H1 (green). *Right panel:* The quantified signal for GST-RhoA G17A associated GEF-H1 from BDP9066 treated cells was reduced by the non-specific GEF-H1 signal associated with GST and the difference was divided by the input GEF-H1 signal, and the resulting ratio was normalized to the equivalent ratio from DMSO treated cells for each independent replicate. Means ± SEM, N = 7. Statistical analysis used an unpaired t-test with Welch’s correction producing the displayed *p* value. The levels of GEF-H1 affinity purified with GST-RhoA G17A after treatment with DMSO vehicle control or 1 µM BDP9066 for 1 h were determined as described above for **C.** A2780 and **D.** OC-1 cells. Means ± SEM, N = 3. Statistical analysis used unpaired t-tests with Welch’s correction producing the displayed *p* values.

### MRCK regulates RhoA activity

Given the observed increased GEF-H1 binding to nucleotide-free RhoA in cells with reduced expression or pharmacological inhibition of MRCK, the possibility of concomitant RhoA activation was examined. Treatment of OVCAR8 cells with BDP9066 from 0.01 to 100 µM for 1 h followed by pulldowns from cell lysates of active GTP-loaded RhoA with the Rhotekin Rho-binding domain (RBD) fused to GST revealed dose-dependent increased RhoA activation at all concentrations, reaching a maximum 3-fold increase and with an EC_50_ = 7.8 µM (**Figures 5A and S4C**). The YAP transcription factor is regulated via phosphorylation on Ser127 by the LATS kinases, which regulates its nuclear-cytoplasmic localization and consequent transcriptional activity ^41^. Inhibition of MRCK activity with BDP9066 was reported to increase YAP phosphorylation in BT549 and HSS78T human breast cancer cells ^19^, while RhoA inhibition with the ADP-ribosyltransferase C3 also increased YAP phosphorylation in HEK293A human embryonic kidney cells ^42^. Treatment of OVCAR8 cells with 3 µM BDP for 16 h significantly (*p* = 0.009) increased YAP phosphorylation by >90%, while treatment with 0.4 µM of cell-permeable FLAG-epitope tagged C3 also significantly (*p* < 0.0001) increased YAP phosphorylation by 4.6-fold (**Figures 5B and S4D**). However, consistent with RhoA activation following BDP9066 treatment, 3 µM BDP9066 significantly (*p* = 0.0042) reduced the level of YAP phosphorylation induced by 0.4 µM FLAG-C3 alone. Active RhoA binds to the SLK kinase to promote phosphorylation on a conserved Threonine residue on Ezrin, Radixin and Moesin (ERM, T567, T564, and T558, respectively) ^43^ to open their conformation and enable coupling of F-actin to the plasma membrane and the promotion of cell retraction ^44^. Treatment of OVCAR8 cells with 3 µM BDP9066 for 16 h did not affect ERM phosphorylation, but 0.4 µM FLAG-C3 significantly (*p* < 0.0001) reduced ERM phosphorylation at the same molecular weight as total Ezrin by 60% (**Figures 5C and S4E**). Consistent with RhoA activation following MRCK inhibition, 3 µM BDP9066 significantly (*p* = 0.0106) reversed the inhibition of ERM phosphorylation induced by 0.4 µM FLAG-C3 alone by ∼50%. The patient-derived OC-1 cells had relatively low levels of ERM phosphorylation, which were increased by treatment with BDP9066 for 1 h in a dose-dependent manner (**Figures 5D and S4F**), paralleling the increased levels of active RhoA observed following BDP9066 treatment of OVCAR8 cells (**Figure 5A**). Collectively, these results directly and indirectly support the conclusion that MRCK represses RhoA activity.

**Figure 5.**
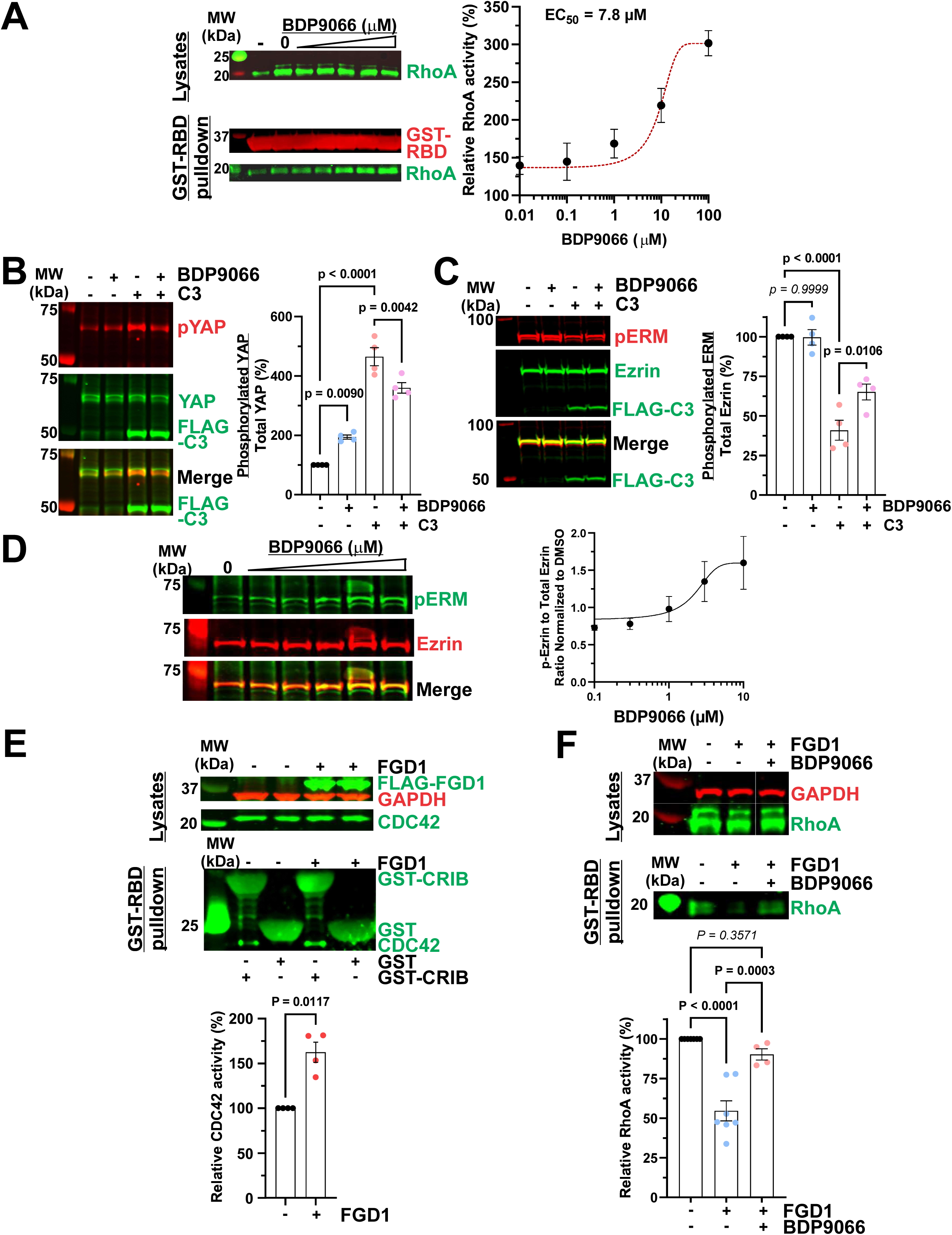
MRCK deletion or inhibition activates RhoA. **A.** *Left panel:* OVCAR8 cells were treated with DMSO vehicle control or indicated BDP9066 concentrations for 1 h, and cell lysates were quantitatively western blotted with antibodies against RhoA (green). Each cell lysate was incubated with GST-RBD protein immobilized on reduced glutathione-conjugated Sepharose beads, and washed glutathione Sepharose beads with immobilized GST-RBD associated with RhoA were quantitatively western blotted with antibodies against GST (red) or RhoA (green). – indicates a serum-starved negative control condition that was not part of the quantitative analysis. *Right panel:* The quantified signal for GST-RBD associated RhoA for each BDP9066 concentration was reduced by its paired non-specific RhoA signal associated with GST and the difference was divided by the corresponding input RhoA signal, and the resulting ratio was normalized to the equivalent ratio from NTC cells for each independent replicate. Means ± SEM, N = 4. Curve-fitting to calculate EC_50_ was by log(drug) versus response with a 4-parameter variable slope. **B.** OVCAR8 cells were treated with DMSO vehicle control, or 3 µM BDP9066 and 0.4 µM cell-permeable FLAG-C3 singly or in combination as indicated for 16 h. Cell lysates were quantitatively western blotted with antibodies against Ser127 phosphorylated YAP (pYAP, red), total YAP (green) or FLAG-epitope tagged C3 (green). The ratio of phosphorylated YAP to total YAP for each condition was normalized to DMSO treated cells for each independent replicate. **C.** OVCAR8 cells were treated with DMSO vehicle control, or 3 µM BDP9066 and 0.4 µM cell-permeable FLAG-C3 singly or in combination as indicated for 16 h. Cell lysates were quantitatively western blotted with antibodies against a phosphorylation on a conserved Threonine residue on Ezrin, Radixin and Moesin (pERM, T567, T564, and T558, respectively, red), total Ezrin (green) or FLAG-epitope tagged C3 (green). The ratio of phosphorylated ERM to total Ezrin for each condition was normalized to DMSO treated cells for each independent replicate. Means ± SEM, N = 4. Statistical analysis used one-way ANOVA with Šídák’s multiple comparisons test between indicated conditions producing the displayed *p* values. **D.** OC-1 cells were treated with DMSO vehicle control or increasing concentrations of BDP9066 from 0.1 to 10 µM for 1 h. Cell lysates were quantitatively western blotted with antibodies against a phosphorylation on a conserved Threonine residue on Ezrin, Radixin and Moesin (pERM, T567, T564, and T558, respectively, green) or total Ezrin (red). The ratio of phosphorylated ERM to total Ezrin for each condition was normalized to DMSO treated cells for each independent replicate. Means ± SEM, N = 3. Curve was generated using log(drug) versus response using four-parameter variable slope nonlinear curve fitting. **E.** OVCAR8 cells transfected with empty vehicle control or the doxycycline-inducible FLAG-tagged FGD1 construct were treated with 1 μg/mL doxycycline for 48 h as indicated, then were lysed and western blotted with antibodies against the FLAG epitope (green), GAPDH (red) and CDC42 (green). Cell lysates, with or without FGD1 expression, were incubated with purified GST or GST-CRIB protein immobilized on reduced glutathione-conjugated Sepharose beads. Washed glutathione Sepharose beads with immobilized GST-CRIB associated with CDC42 were quantitatively western blotted with antibodies against CDC42 (green). The quantified signal for GST-CRIB associated CDC42 for each condition was reduced by the non-specific CDC42 signal associated with GST and the difference was divided by the input CDC42 signal, and the resulting ratio was normalized to the equivalent ratio from untreated cells for each independent replicate. Means ± SEM, N = 4. Statistical analysis used unpaired t-tests with Welch’s correction producing the displayed *p* values. **F.** OVCAR8 cells expressing doxycycline-inducible FLAG-tagged FGD1 were treated with 1 μg/mL doxycycline for 48 h and 3 µM BDP9066 for 1 h as indicated. Cell lysates, with or without FGD1 expression and BDP9066 treatment, were incubated with purified GST or GST-RBD protein immobilized on reduced glutathione-conjugated Sepharose beads. Washed glutathione Sepharose beads with immobilized GST-RBD associated with RhoA were quantitatively western blotted with antibodies against RhoA (green). The quantified signal for GST-RBD associated RhoA for each condition was reduced by the non-specific RhoA signal associated with GST and the difference was divided by the input RhoA signal, and the resulting ratio was normalized to the equivalent ratio from untreated cells for each independent replicate. Means ± SEM, N = 4. Statistical analysis used one-way ANOVA with Tukey’s multiple comparison test between indicated conditions producing the displayed *p* values.

To activate endogenous MRCK, OVCAR8 cells were stably transfected to express a FLAG-tagged constitutively active version of the CDC42-specific GEF Faciogenital Dysplasia 1 (FGD1) composed of the catalytic Dbl Homology (DH) and Pleckstrin Homology (PH) domains (amino acids 375-710) in a doxycycline-inducible manner. Titration with doxycycline revealed efficient FGD1 induction at all concentrations from 0.125 μg/mL (**Figure S5A**). Treatment of cells transfected with an empty vector control or with the FLAG-tagged FGD1 construct with 1 μg/mL doxycycline for 48 h followed by pulldowns from cell lysates of active GTP-loaded CDC42 with the PAK1 CDC42/Rac Interacting-Binding domain (CRIB) fused to GST revealed significant (*p* = 0.0117) > 60% increased activation of CDC42 in FGD1 expressing cells (**Figure 5E and S5B**). In parallel, the induction of FGD1 expression significantly (*p* < 0.0001) decreased RhoA activity by ∼45%, which could be reversed by 1 h treatment with 3 µM BDP9066 (**Figure 5F and S5C**).

Transient GEF-H1 knockdowns were used to determine if the observed RhoA activation in cells following pharmacological MRCK inhibition or MRCK knockdown were dependent on GEF-H1 expression. GEF-H1 knockdown with two independent shRNA constructs reversed the significant (*p* = 0.0132) 90% increase in RhoA activity in cells treated 10 µM BDP9066 for 1 h relative to non-targeted control cells (**Figures 6A-B and S6A**). In addition, siRNA-mediated GEF-H1 knockdown in OC-1 cells reversed BDP9066 induced RhoA activation, as well as the increased RhoA activation observed after shRNA-mediated MRCKα/MRCKβ knockdown (**Figure S6B**). Collectively, these results demonstrate that MRCK regulates RhoA activity via GEF-H1.

**Figure 6.**
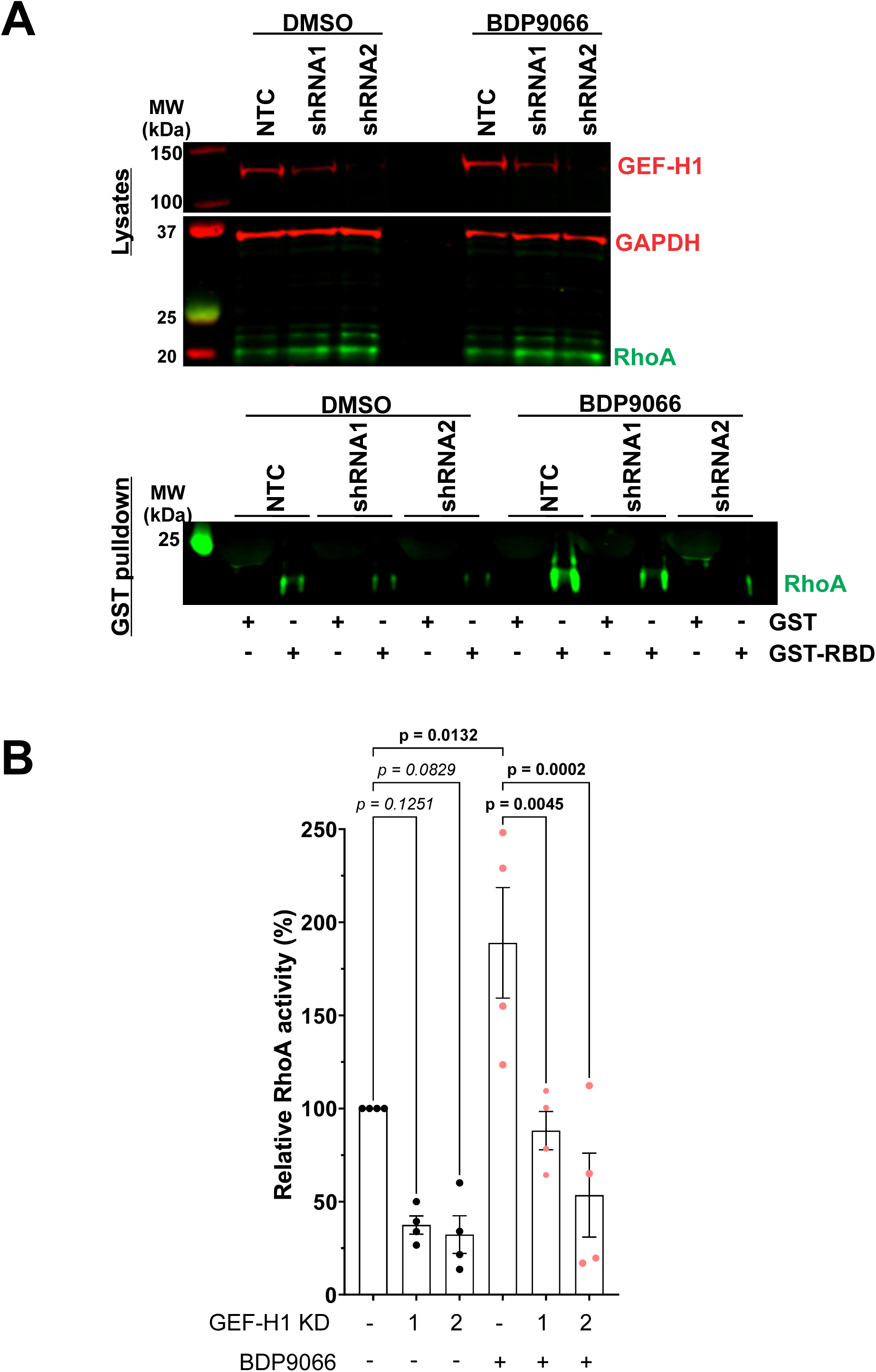
RhoA activation induced by MRCK deletion or inhibition is GEF-H1 dependent. **A.** OVCAR8 cells were transfected with plasmids encoding doxycycline-inducible non-targeted control (NTC) shRNA or GEF-H1 targeted shRNAs (shRNA1, shRNA2). After doxycycline treatment for 48 h, cells were treated with DMSO vehicle control or 10 µM BDP9066 for 1 h, and cell lysates were western blotted with antibodies against GEF-H1 (red), GAPDH (red) and RhoA (green). Cell lysates, with or without GEF-H1 knockdown (KD), were incubated with purified GST or GST-RBD protein immobilized on reduced glutathione-conjugated Sepharose beads. Washed glutathione Sepharose beads with immobilized GST-RBD associated with RhoA were quantitatively western blotted with antibodies against RhoA (green). **B.** The quantified signal for GST-RBD associated RhoA for each condition was reduced by the non-specific RhoA signal associated with GST and the difference was divided by the input RhoA signal, and the resulting ratio was normalized to the equivalent ratio from DMSO treated NTC cells for each independent replicate. Means ± SEM, N = 4. Statistical analysis used one-way ANOVA with Tukey’s multiple comparison test between indicated conditions producing the displayed *p* values.

### MRCK phosphorylates GEF-H1

To determine if GEF-H1 is a potential MRCKα substrate, we performed *in vitro* kinase assays using bacterially-expressed GST-GEF-H1 and recombinant MRCKα kinase domain (1-473 aa) or full-length FLAG-MRCKα purified from HEK293 cells, and measured phosphorylation using LC-MS/MS. Full-length and kinase domain MRCKα phosphorylated several Serine residues within GST-GEF-H1, including S121, S122, S151, S174 and S886 (**Figures 7A, S7A-I, Data File S2**), with S151 phosphorylation by FLAG-MRCKα exhibiting the greatest LFQ intensity followed by S174 (**Figures S7A**). Notably, several phosphorylation sites are located in an N-terminal GEF-H1 region near the catalytic Dbl Homology domain (DH), that was determined to bind MRCKα (**Figure 3B**) and which has been shown to bind to the dynein light chain protein Tctex-1 to repress GEF-H1 RhoGEF activity ^37^. Mapping of the identified GEF-H1 phosphorylation sites using AlphaFold2 ^45,46^ revealed their location on disordered loop regions of the protein (**Figure 7B**).

**Figure 7.**
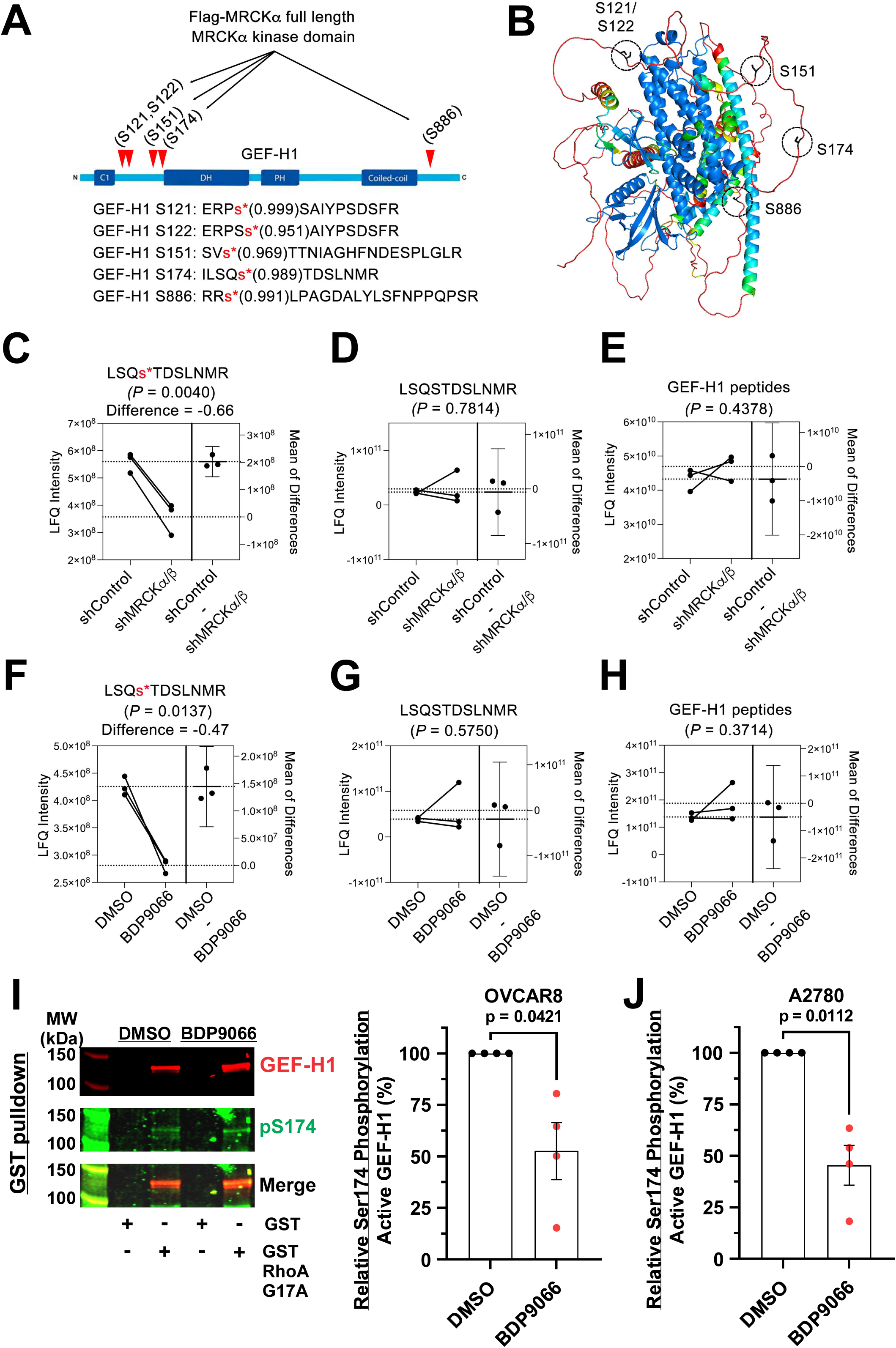
MRCK phosphorylation of GEF-H1 on Serine 174. **A.** GST-GEF-H1 tryptic peptides phosphorylated by full length FLAG-MRCKα and/or MRCKα kinase domain (1-473 aa) determined by *in vitro* kinase assay. The phosphorylated residue is denoted by s* and the localization probability of phosphorylation determined by MaxQuant B. AlphaFold2 model of predicted MRCKα phosphorylation sites within GEF-H1. Estimation plots depicting LFQ intensity and Mean of Differences of **C.** S174 phosphorylated, **D.** S174 non-phosphorylated or **E.** total GEF-H1 peptides from FLAG-GEF-H1 expressing OC-1 cells following 96 hours of MRCKα/β knockdown. FLAG-GEF-H1 was immunoprecipitated using anti-FLAG antibodies, trypsin digested, and peptides analysed by LC-MS. Differences in log_2_ LFQ intensities among FLAG-GEF-H1 tryptic peptides were determined by paired t test with adjusted *p* value cutoff of 0.05 using Perseus Software. Mean of differences of paired t test analysis was performed in Prism Software. Estimation plots depicting LFQ intensity and Mean of Differences of **F.** S174 phosphorylated, **G.** S174 non-phosphorylated or **H.** total GEF-H1 peptides from FLAG-GEF-H1 expressing OC-1 cells following 24 h treatment with 5 µM BDP9066. FLAG-GEF-H1 was immunoprecipitated using anti-FLAG antibodies, trypsin digested, and peptides analysed by LC-MS. Differences in log_2_ LFQ intensities among FLAG-GEF-H1 tryptic peptides were determined by paired t test with adjusted *p* value cutoff of 0.05 using Perseus Software. Mean of differences of paired t test analysis was performed in Prism Software. **I.** *Left panel:* OVCAR8 cells were treated with DMSO vehicle control or 10 µM BDP9066 for 1 h, as indicated. DMSO or BDP9066 treated cell lysates were incubated with purified GST or nucleotide free mutant GST-RhoA G17A protein immobilized on reduced glutathione-conjugated Sepharose beads as indicated. Washed glutathione Sepharose beads with immobilized GST fusion proteins associated with GEF-H1 were quantitatively western blotted with antibodies against GEF-H1 (red) and pS174 (green). *Right panel:* The ratios of pS174 to total GEF-H1 associated with nucleotide-free GST-RhoA G17A for BDP9066 treated OVCAR8 cells were normalized to DMSO treated cells for each independent replicate. **J.** The proportion of pS174 to total GEF-H1 associated with nucleotide-free GST-RhoA G17A for BDP9066 treated A2780 cells was determined as described above. Means ± SEM, N = 4. Statistical analysis used an unpaired t-tests with Welch’s correction producing the displayed *p* values.

We next determined whether the *in vitro* MRCKα phosphorylation sites on GEF-H1 were dependent on MRCKα/β activity in OC-1 cells. We expressed FLAG-GEF-H1 in OC-1 cells depleted of MRCKα/β via shRNA mediated knockdown, immunoprecipitated FLAG-GEF-H1 and measured GEF-H1 phosphorylation using LC-MS/MS. Notably, the phosphorylated S174 tryptic peptide LFQ intensity was reduced in FLAG-GEF-H1 expressing OC-1 cells following MRCKα/β knockdown compared to control (Log_2_-fold difference = -0.66, *p* = 0.0040) (**Figure 7C, Data File S3**), while there were no significant differences in LFQ intensity of the non-phosphorylated S174 (*p* = 0.7814) (**Figure 7D**) or total GEF-H1 (*p* = 0.4378) peptides (**Figure 7E**) observed following MRCKα/β knockdown. In addition, there were no significant reductions observed in LFQ intensities of phosphorylated S122, S151 or S886 tryptic peptides in response to MRCKα/β knockdown in OC-1 cells (**Figure S7J**). Treatment of FLAG-GEF-H1 expressing OC-1 cells with 5 µM BDP9066 for 24 h significantly reduced the LFQ intensity of the phosphorylated S174 tryptic peptide compared to DMSO (Log_2_-fold difference = -0.47, *p* = 0.0137) (**Figure 7F, Figure S7K, Data File S4**), while no significant changes in LFQ intensities of the non-phosphorylated S174 or total GEF-H1 peptides were observed (**Figures 7G-H**). Taken together, the mass spectrometry findings demonstrated that the GEF-H1 S174 phosphorylation site was sensitive to both depletion of MRCKα/β protein and pharmacological kinase inhibition.

A phospho-selective antibody was developed to detect GEF-H1 S174 phosphorylation (pS174). The pS174 signal on GEF-tagged GEF-H1 expressed in HEK293T cells was significantly (*p* < 0.0001) reduced by ∼50% in cells treated with 10 µM BDP9066 for 1 h, and by ∼80% when the S174 was mutated to Alanine (S174A), validating the antibody for detection of GEF-H1 S174 phosphorylation (**Figure S8A**). In contrast to the reduction in S174 phosphorylation induced by BDP9066, treatment of OVCAR8 cells with 10 µM BDP9066 for 1 h did not affect the phosphorylation of Ser886 as determined by western blotting (pS886; **Figure S8B**), a site previously shown to be phosphorylated by kinases including the CDC42-regulated PAK1 and PAK4, protein kinase A, Aurora A and PAR1b ^37^. To determine how the GEF-H1 activation state corresponded to S174 phosphorylation, OVCAR8 cells were treated for 1 h with DMSO or 10 µM BDP9066 and active GEF-H1 was affinity purified with nucleotide-free GST-RhoA G17A. As seen previously (**Figure 4B**), BDP9066 treatment increased the amount of GEF-H1 associated with nucleotide-free GST-RhoA G17A relative to DMSO treatment, but with significantly (*p* = 0.0421) ∼50% proportionally decreased S174 phosphorylation (**Figures 7I and S8C**). Similarly, BDP9066 treatment significantly (*p* = 0.0112) decreased the proportion of S174 phosphorylated GEF-H1 that had been affinity purified with RhoA G17A by ∼55% in A2780 cells (**Figure 7J**). Collectively, these observations indicate that MRCKα phosphorylates S174 on GEF-H1, which is paralleled by increased GEF-H1 association with nucleotide-free RhoA.

### MRCK influences GEF-H1 association with tubulin

The primary mechanism of GEF-H1 regulation is mediated through its association with microtubules, which can occur directly through its N- and C-termini ^47,48^ and its pleckstrin homology (PH) domain ^49^. GEF-H1 can also bind to other proteins, such as the dynein motor light-chain Tctex-1 ^50^, to facilitate association with microtubules. To determine if MRCK influenced GEF-H1 association with microtubules, OVCAR8 cells were treated with 10 µM BDP9066 or DMSO for 1 h, followed by immunoprecipitation with α-Tubulin antibodies. While BDP9066 treatment did not affect the ability of α-Tubulin to be immunoprecipitated, there was a significant (*p* = 0.0455) ∼40% decrease in co-immunoprecipitated GEF-H1 (**Figures 8A and S9**). There was a similar 34% decrease in GEF-H1 that co-immunoprecipitated from BDP9066 treated A2780 cells (**Figure 8B**). Taken together, these observations indicated that there was reduced association between GEF-H1 and microtubules when MRCK was inhibited, consistent with consequent increased RhoA activation (**Figure 6**).

**Figure 8.**
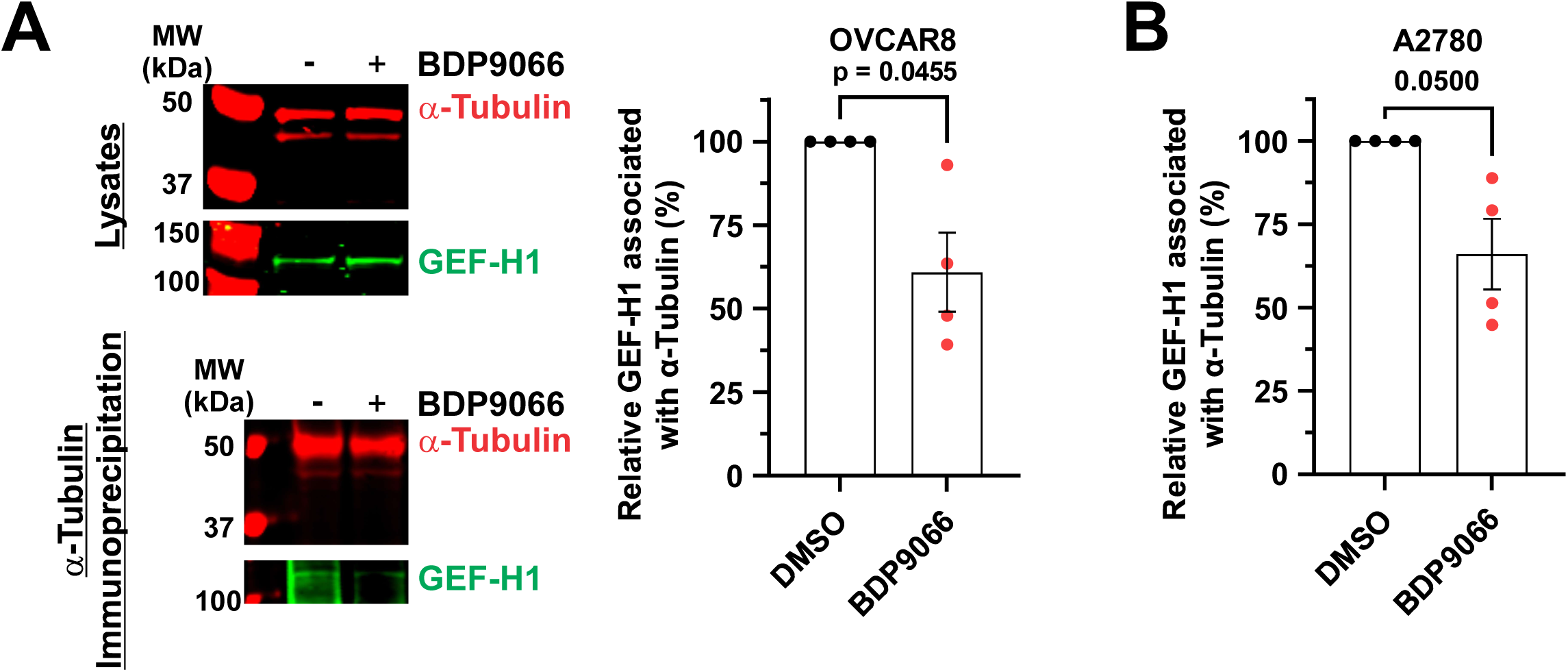
MRCK inhibition reduces GEF-H1 association with α-Tubulin. **A.** *Left panel:* OVCAR8 cells were treated with DMSO vehicle control (-) or 10 µM BDP9066 (+) for 1 h, as indicated. Cell lysates were quantitatively western blotted with antibodies against α-Tubulin (red) and GEF-H1 (green). Cell lysates were incubated with antibodies against α-Tubulin, and immunoprecipitated proteins were quantitatively western blotted with antibodies against α-Tubulin (red) and GEF-H1 (green). *Right panel:* The ratios of GEF-H1 to total α-Tubulin for BDP9066 treated OVCAR8 cells were normalized to the input levels of each protein and then expressed relative to DMSO treated cells for each independent replicate. **B.** The amount of GEF-H1 associated with immunoprecipitated α-Tubulin from A2780 cells treated with DMSO vehicle control or 10 µM BDP9066 for 1 h was determined as described above. Means ± SEM, N = 4. Statistical analysis used an unpaired t-tests with Welch’s correction producing the displayed *p* values.

### MRCK inhibition of GEF-H1 facilitates cell-cell interactions

Interactions between cells that lead to cell-cell adhesions to be formed require a balance between propulsive forces that enable cell-to-cell contacts to be established versus contractile forces that power cell retraction ^51^. OVCAR8 cells are epithelial-like, based on their morphology, ability to form 3D spheroids and expression patterns of epithelial and mesenchymal markers ^52^. Live cell timelapse imaging of OVCAR8 cells expressing LifeAct-TagRFP-T and GFP-RhoA enabled the identification of cell pairs moving towards each other to increase the extent of their interactions. The addition of 3 µM BDP9066 to control cells resulted in rapid retraction of each paired cell from the other (**Figure 9A** left panels; **Video 1**). In contrast, GEF-H1 knockdown markedly attenuated the retraction induced by BDP9066 (**Figure 9A** right panels; **Video 2**). To quantify this effect, the change in position of the centroid of the nuclei in each GFP-RhoA positive cell pair was measured relative to their position at the time of DMSO or BDP9066 addition (t = 0). BDP9066 treatment of cells transfected with non-targeting control siRNA led to significant (*p* < 0.0001) movement of cells away from each other relative to non-targeting control siRNA transfected cells treated with DMSO (**Figure 9B**). Furthermore, the movement of nuclei away from each other induced by BDP9066 in control cells was significantly (*p* < 0.0001) reduced when GEF-H1 was knocked down (**Figure 9B**), Although NTC siRNA transfected cell pairs treated with DMSO vehicle control appeared to trend towards continued movement together, there was no significant difference (*p* = 0.2593) compared to BDP9066 treated cells in which GEF-H1 was knocked down (**Figure 9B, Video 3**).

**Figure 9.**
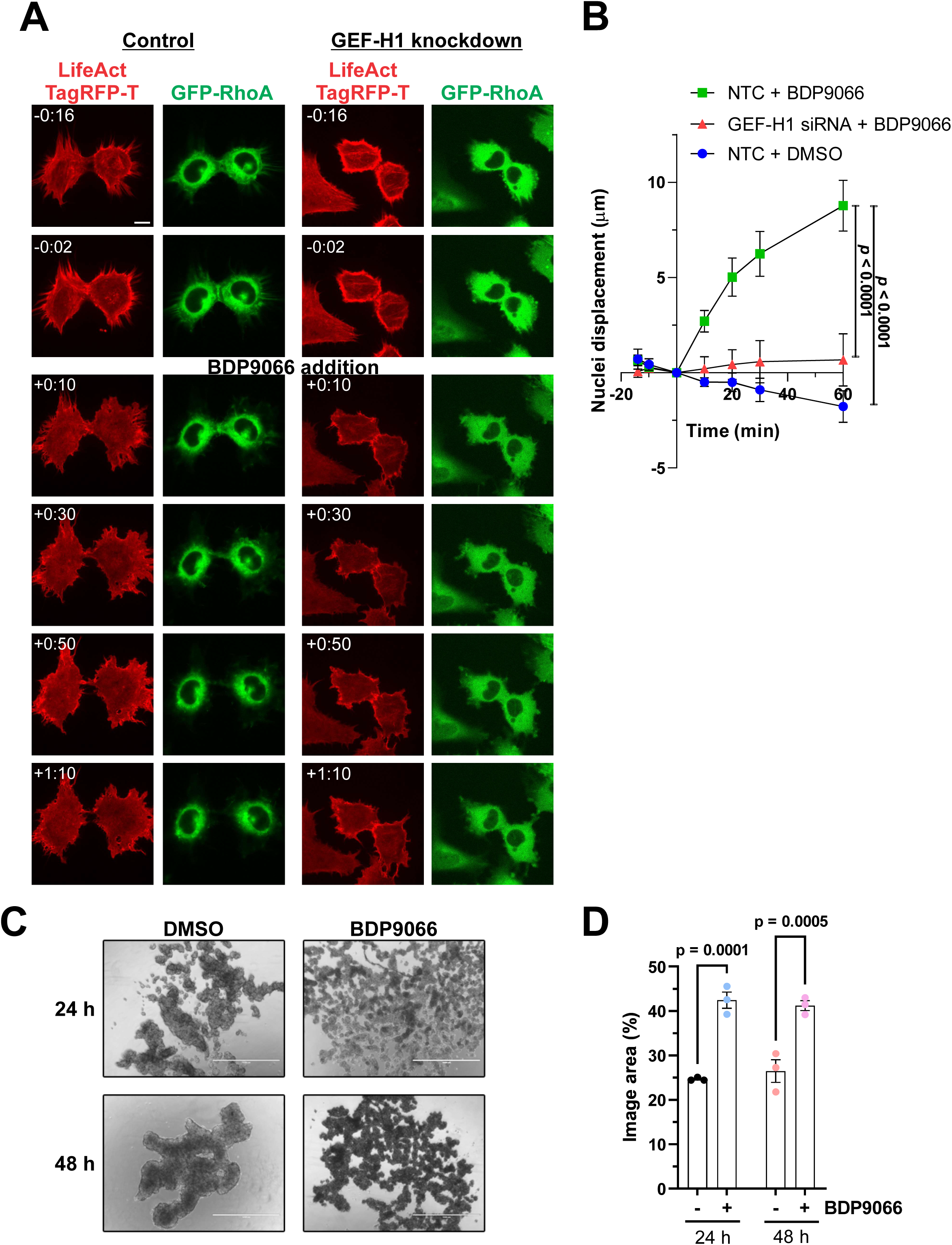
Cell retraction induced by MRCK inhibition is reduced by GEF-H1 knockdown. **A.** Live OVCAR8 cells stably expressing LifeAct-TagRFP and GFP-RhoA, which were either untransfected controls or transfected with siRNA to knockdown GEF-H1, as indicated, were imaged by spinning disk confocal microscopy before and after addition of 3 µM BDP9066. Representative images at indicated time points were acquired from timelapse videos. Scale bar = 10 µm. **B.** The movement of nuclei centroids in GFP-RhoA images relative to their positions at t = 0 min (DMSO or BDP9066 addition). Positive values indicate movement away from their t = 0 position relative to its cell partner, negative values indicate movement towards its cell partner. Means ± SEM. N = 13 for non-targeting control (NTC) transfected cell pairs treated with DMSO, N = 18 for NTC transfected cell pairs treated with BDP9066, N = 12 for GEF-H1 siRNA transfected cell pairs treated with BDP9066. Statistical analysis used two-way ANOVA with Tukey’s multiple comparisons test between indicated conditions producing the displayed *p* values. **C.** Equal numbers of OVCAR8 cells were plated in low adhesion tissue culture plates in the presence of DMSO or 5 µM BDP9066, and imaged 24 h and 48 h later. Scale bar = 400 µm. **D.** Percentage areas occupied by cells were determined for each image. Means ± SEM. N = 3 for each condition. Statistical analysis used one-way ANOVA with Šídák’s multiple comparisons test between indicated conditions producing the displayed *p* values.

To determine if the effects of BDP9066 on cell-cell contacts would also affect spheroid formation, equal numbers of OVCAR8 cells were plated as single cell suspensions onto low adhesion tissue culture plates with either DMSO vehicle control or 5 µM BDP9066. Imaging wells 24 h and 48 h after plating revealed that OVCAR8 cells in DMSO were able to form large aggregates that became more compacted over time (**Figure 9C**, left panels). In contrast, BDP9066 treated OVCAR8 cells were impaired in their ability to effectively associate and compact (**Figure 9C**, right panels). Quantification of the percentage area of each image occupied by cells showed that BDP9066 treated cells filled significantly more area at 24 h (*p* = 0.001, 72% larger) and 48 h (*p* = 0.0005, 55% larger) (**Figure 9D**). These results support the conclusion that the MRCK-induced repression of GEF-H1 mediated RhoA activation enables the formation of cell-cell contacts to contribute to the formation of compacted 3D HGSOC multicellular structures.

To determine if the effects of MRCK inhibition on spheroid formation would impact cell viability, four human HGSOC patient-derived organoid (PDO) models were tested for their sensitivity to the MRCK inhibitor BDP9066. PDOs are self-organizing microtissues grown within 3D extracellular matrices that are valuable preclinical models for studying cancer with significant advantages over other model systems ^53^. Numerous studies have determined that PDO drug sensitivity profiles are significantly correlated with patient responses, making them valuable pre-clinical surrogates ^54^. PDOs were dissociated to single cells and seeded on Matrigel basement membrane matrix, then after 24 h were treated with 0.01 to 50 µM BDP9066, or DMSO vehicle control, for 6 days. Cell viability assays revealed that all 4 PDOs were sensitive to BDP9066 with GI_50_ values ranging from 1.36 to 5.29 µM (**Figure 10A**). As a control for this effect being due to on-target MRCK inhibition and not the chemical composition of the inhibitor, the active enantiomer BDP9066 (K_i_ = 0.02332 nM for MRCKβ at 100 µM ATP) and the chemically identical but 67-fold less active enantiomer BDP9069 (K_i_ = 1.563 nM for MRCKβ at 100 µM ATP) ^10^ were tested on the OPTO.129 PDO model. While BDP9066 produced a dose dependent reduction in cell viability (GI_50_ = 2.35 µM), there was less than 25% reduction in cell viability at all concentrations up to 50 µM of BDP9069 (**Figure S10**). In comparison to the effect of BDP9066 on PDO viability, the Food and Drug Administration and Health Canada approved ovarian cancer therapeutic carboplatin had an average GI_50_ (16.7 µM) that was 5.2 fold higher than the average GI_50_ (3.2 µM) for BDP9066 for the same 3 PDOs (**Figure 10B**), with statistically significant (*p* = 0.0138) differences between the pairwise sensitivity of each PDO for BDP9066 versus carboplatin (**Figure 10C**). Taken together, these findings support a model in which MRCKα promotes HGSOC spheroid formation and growth *via* the negative regulation of GEF-H1 mediated RhoA activation (**Figures 10D-E**)

**Figure 10.**
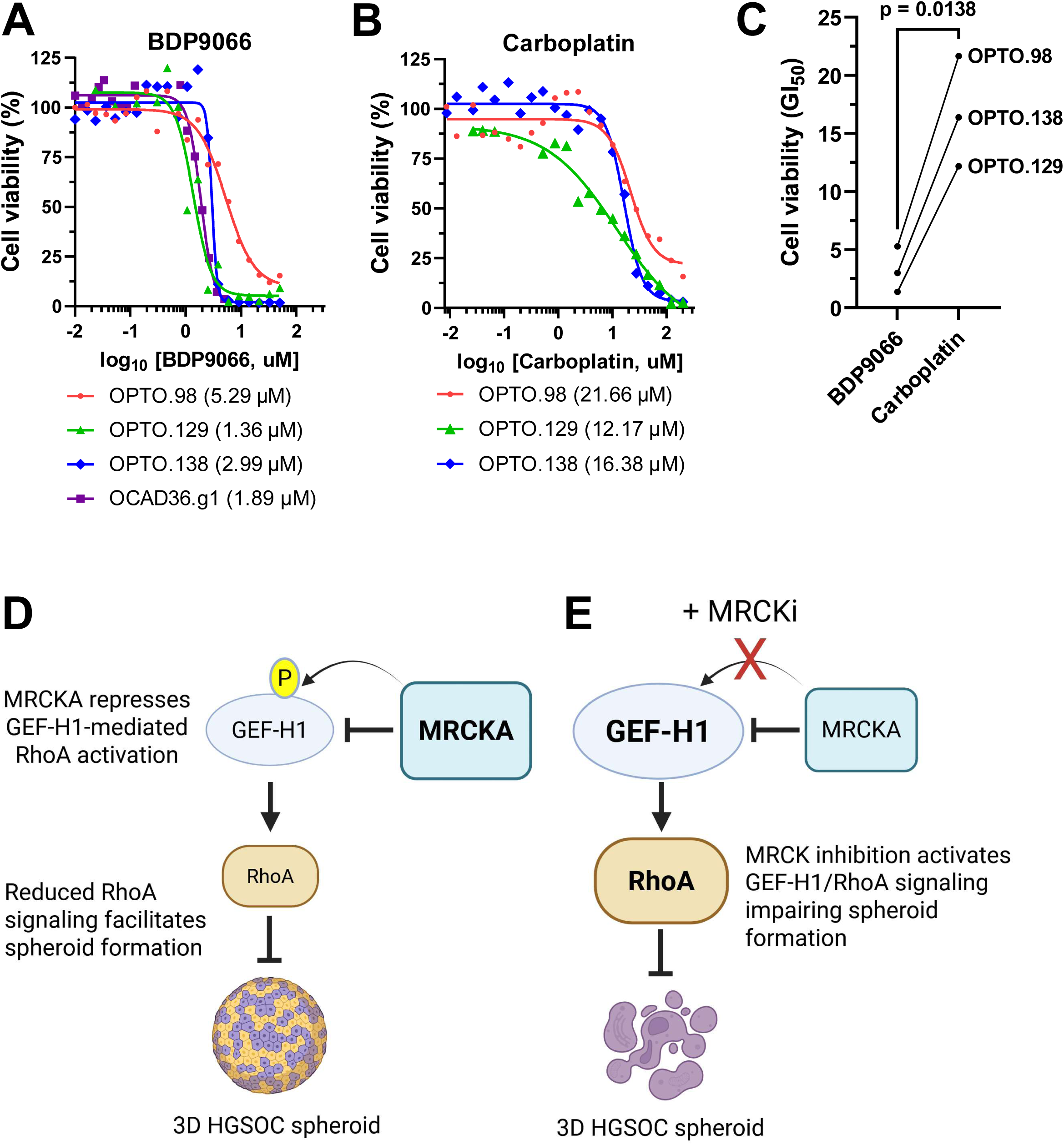
MRCK inhibition decreases patient-derived organoid viability. HGSOC patient derived organoids grown on Matrigel were treated with DMSO vehicle control or indicated concentrations of **A.** BDP9066 or **B.** Carboplatin for 6 days. Relative GI_50_ values were calculated from 21-point drug concentrations of log(drug) versus response using four-parameter variable slope nonlinear curve fitting. **C.** Calculated GI_50_ values for BDP9066 and Carboplatin were plotted for each PDO. Statistical analysis used a paired t-test producing the displayed *p* value. **D.** Model of MRCKα promotion of HGSOC spheroid growth via negative regulation of GEF-H1-mediated RhoA activation. GEF-H1 activates RhoA reducing cell-cell contact and spheroid formation. MRCKα represses GEF-H1-mediated activation of RhoA via protein interaction and/or phosphorylation facilitating HGSOC spheroid growth. **E.** Blockade of MRCK activity alleviates this repression resulting in the activation of GEF-H1 and RhoA signaling that impairs HGSOC spheroid formation, growth and survival.

## Discussion

The dynamic organization of the actin cytoskeleton responds to changes in the activation status of the associated myosin complex ^55^. One of the most important events regulating the actin-myosin cytoskeleton is phosphorylation of the myosin II regulatory light chains (MLC), part of the “conventional” and widely-expressed myosin II complex ^55^. MLC phosphorylation on two sites, Thr18 and Ser19, induces conformational changes that promote the association of the myosin complex with filamentous actin (F-actin) and increased myosin ATPase activity to provide energy for the “power stroke” that drives actin-myosin contraction ^56^. There are two sets of related kinases that mediate MLC phosphorylation downstream of Rho GTPase activation: the RhoA-regulated ROCK kinases ^21^ and the CDC42-regulated MRCK kinases ^11,12^. Previous research showed that it was necessary to block both ROCK and MRCK pathways to achieve maximal inhibition of MLC phosphorylation and consequent reduction of tumor cell invasion into 3D matrices ^57,58^, which could be interpreted as being due to additive phosphorylation of common protein substrates including MLC. We now show that that the relationship between CDC42 and RhoA signaling is more complicated, with the CDC42 effector MRCKα directly binding and phosphorylating the RhoA selective guanine nucleotide exchange factor GEF-H1 to regulate its activity. In contrast to the apparent additivity of MRCK and ROCK in mediating MLC phosphorylation ^57,58^, our results revealed that the effects of RhoA inhibition on increasing inactivating YAP phosphorylation (**Figure 5C**) and decreasing activating ERM phosphorylation (**Figure 5D**) were antagonized by MRCK inhibition, indicating that the MRCK and RhoA worked in opposition to regulate these biological responses. Thus, the effect of MRCK inhibition on the phosphorylation of common substrates such as MLC could be partially masked by consequent activation of the GEF-H1/RhoA/ROCK pathway, necessitating combined MRCK plus ROCK inhibition to achieve maximal reductions in substrate phosphorylation.

The spread of ovarian tumor cells differs from other epithelial cancers. Rather than invading through adjacent tissue and intravasating into lymphatic or blood vessels, ovarian cancers detach from primary tumors and form multicellular clusters (spheroids) that disseminate and colonize within the peritoneal cavity ^59^. Late stage HGSOC patients frequently present with fluid build-up within the peritoneal cavity containing numerous tumor cell spheroids (malignant ascites) that facilitate metastasis to additional extraperitoneal sites (*e.g.* lung, liver) ^60^. Spheroids form via a series of distinct steps, beginning with the establishment of cell-cell contacts that enable the formation of loose aggregates, followed by compaction to form more tightly associated cell clusters ^23^. The Rho GTPase regulated actin cytoskeleton makes essential contributions at each step, necessitating a switch between Rac1 and CDC42 activation during the early stages to enable the formation of cell-cell contacts ^24^ and subsequent RhoA activation ^28^ to promote actin-myosin contraction to drive spheroid compaction^29^. We now show that the CDC42-associated MRCKα kinase facilitates the formation of cell-cell contacts between HGSOC cells *via* its ability to regulate the RhoA selective GEF-H1 protein (**Figures 10D-E**). While pharmacological MRCK inhibition of cell pairs resulted in rapid retraction from cell-cell contacts, this response was effectively blocked by GEF-H1 knockdown (**Figure 9**), thus allowing cells to maintain their interactions. Given that contact inhibition of locomotion (CIL) typically results in RhoA activation at cell-cell contact sites to collapse protrusions and promote re-polarization to enable migration of each cell in the opposite direction ^61^, one interpretation is that MRCKα acts to antagonize CIL by inhibiting GEF-H1 and RhoA activation to promote cell-cell interactions. Pharmacological MRCK inhibition with BDP9066 rapidly resulted in movement of cells away from each other (**Figures 9A-B**) and reduced their ability to form aggregates (**Figures 9C-D**), thus resembling activation of CIL. Inhibition of CIL has also been shown to contribute to heterotypic cell-cell interactions that promote invasion and metastasis ^62^, consistent with a role for increased MRCK signaling in metastatic breast ^17^ and ovarian cancers ^9^.

Multicellular HGSOC spheroids spread within the peritoneal cavity, aided by the flow of the fluidic component of malignant ascites ^63^. Clustering into 3D spheroids increases the resistance of HGSOC cells to anoikis, thus facilitating proliferation and spread in suspension and in the absence of survival signals that would typically be generated by attachment to extracellular matrix ^64,65^. By antagonizing the formation of cell-cell contacts necessary for the establishment of spheroids, MRCK inhibition would remove the anti-anoikis survival advantage conferred by cell clustering. Consistent with this conclusion, the viability of patient-derived HGSOC cells was significantly diminished by pharmacological MRCK inhibition in a dose-dependent manner (**Figure 10A**).

The establishment of organoids from ovarian cancers makes use of small molecule ROCK inhibition for initiation and passaging after dissociation ^66^. Furthermore, organoids derived from a variety of tissues including intestine, liver, pancreas, lung and brain are routinely cultured in the presence of small molecule ROCK inhibitors to promote the survival of stem/progenitor cells, facilitate cell aggregation and reduce anoikis ^67^. This general requirement to inhibit ROCK activity to enable the formation of 3D multicellular clusters supports the concept that there would be a distinct advantage to increase MRCKα expression or activity in order to limit RhoA/ROCK signalling, thus contributing to the formation and growth of ovarian cancer spheroids.

## Methods

Details of antibodies, plasmids, siRNA/shRNA/sgRNA, buffers and purification materials are provided in the Supplemental Methods spreadsheet.

### Cell culture and drug treatments

Human-embryonic kidney HEK293T and HEK293FT, human epithelial ovarian carcinoma OVCAR8, human ovarian endometroid adenocarcinoma A2780, human ovarian adenocarcinoma SK-OV-3, and patient-derived high-grade serous ovarian carcinoma OC-1 cells were cultured and maintained in a humidified atmosphere at 37°C and 5% CO_2_. OVCAR8, A2780, and SK-OV-3 cell lines were cultured in RPMI 1640 Medium (Sigma Aldrich) supplemented with 10% fetal bovine serum (FBS – Gibco) and 100 units/mL of penicillin-streptomycin (P/S – Gibco), while OC-1 cell media was further supplemented with 2 mM L-Glutamine (BioShop) and 5 µg/mL of insulin (Sigma Aldrich). HEK293T and HEK293FT cells were cultured in DMEM (Sigma Aldrich), supplemented with 10% FBS and 100 units/mL P/S. All cell lines were passaged using 0.05% Trypsin/EDTA (Gibco). Cell lines were passaged 24-48 hours prior to drug treatment. Once cells reached 60-80% confluency, cell media was replaced with complete media supplemented with indicated drug concentrations or 0.1% (v/v) DMSO and incubated for indicated times.

### siRNA transfection

Forward Transfection: Cells were transfected using Lipofectamine RNAiMAX Transfection Reagent (Invitrogen) according to manufacturer’s instructions. Cells were cultured to 60-80% confluency in 6 well dishes, 30 pmol of GEF-H1 siRNA (siGEF-H1 1: Dharmacon, D-009883-01-002 ARHGEF2, siGEF-H1 2: Dharmacon, D-009883-02-0010 ARHGEF2, or siGEF-H1 5: Dharmacon, D-009883-05-0010 ARHGEF2), or nonspecific control siRNA (Dharmacon, D-001206-14-05) was diluted in 500 μL Opti-MEM Reduced Serum Medium 1x (Gibco) and combined with 7.5 μL Lipofectamine RNAiMAX reagent diluted in Opti-MEM media. The mixture was incubated for 10-15 minutes at room temperature, then dropped onto cells. After 5 h, media was replaced with complete RPMI (supplemented with 10% FBS and 100 units/mL P/S). Cell lysates were prepared 48 h after transfection.

Reverse Transfection: Cells were passaged into 6-well dishes and transfected on the same day with 30-75 pmol GEF-H1 siRNA 1 GEF-H1 siRNA 2, GEF-H1 siRNA 5, or nonspecific control siRNA using Lipofectamine RNAiMAX reagent, according to the manufacturer’s protocol. siRNAs were diluted in 500 μL Opti-MEM media with 7.5 μL Lipofectamine RNAiMAX and incubated at room temperature for 15 minutes. The siRNA-lipid complexes were then combined with 250,000-375,000 cells detached with 0.05% Trypsin/EDTA and suspended in OptiMEM. Cell lysates were prepared 48 h after transfection.

### Transient transfections

HEK293T cells were grown on 100 mm tissue culture dishes and transfected at ∼ 70% confluency. Cell media was replaced with serum-free DMEM 30 minutes prior to transfection. Cells were transfected with 5 μg plasmid DNA in 500 μL serum-free DMEM combined with a transfection mixture containing 15 μL PolyJet *In Vitro* DNA Transfection Reagent (SignaGen) reagent diluted separately in 500 μL serum-free DMEM. Transfection complexes were incubated for 10-15 minutes at room temperature prior to being added to cells. After 4 h, media was replaced with complete DMEM with 10% FBS and 100 units/mL P/S. Cell lysates were prepared 48 h after transfection.

OC-1 cells were grown on 10 mm tissue culture dishes and transfected at ∼70% confluency. 5 μg of Plasmid DNA was diluted in 500 μL Opti-MEM with 10 μL Lipofectamine PLU reagent (Invitrogen) then combined with 25 μL Lipofectamine LTX (Invitrogen) reagent separately diluted in 500 μL Opti-MEM. Complexes were incubated for 5 minutes at room temperature prior to being added to cells. Lysates were prepared 48 h after transfection.

### Stable transfections

OVCAR8 cells were grown in 6 well dishes and transfected at ∼ 70% confluency. 4 μg of pcDNA 3.1-hygro-LifeAct-TagRFPt Plasmid DNA was diluted in 250 μL Opti-MEM with 10 μL Lipofectamine 2000 reagent (Invitrogen) then combined with 25 μL Lipofectamine LTX reagent separately diluted in 250 μL Opti-MEM. Complexes were incubated for 5 minutes at room temperature prior to being added to cells. Cells were split into 10 cm dishes, and the appropriate selection antibiotic was added when cells reached ∼ 70% confluency, with drug selection continuing for 14 days.

### Lentivirus packaging

HEK293FT cells were grown to 70% confluence in 6 well dishes. At ∼ 70% confluence, media was changed to DMEM supplemented with 100 units/mL P/S and 5% heat inactivated FBS. Cells were transfected with a DNA mixture (1 µg lentivirus plasmid, 1 µg psPAX2 plasmid and 0.5 µg pCMV-VSV-G plasmid in 100 µL DMEM) combined with a transfection mixture containing 6 µL of PolyJet (FroggaBio) in 100 µL DMEM. After 4 hours, cells were incubated in DMEM supplemented with 100 units/mL P/S and 5% heat inactivated FBS for 48 h. Conditioned media containing packaged lentivirus was removed and filtered with 0.45 μM filters (FroggaBio).

### Lentiviral transduction

OVCAR8 or OC-1 cells were grown to ∼ 70% confluency, and media was replaced with filtered conditioned media containing packaged lentivirus for 24 h. Cells were washed 3X with Dulbecco’s Phosphate-Buffered Saline (D-PBS – Sigma-Aldrich), and RPMI 1640 with 10% FBS and 100 units/mL P/S was placed on cells for 24 h. Cells were split into 10 cm dishes, and the appropriate selection antibiotic was added when cells reached ∼ 70% confluency, with drug selection continuing for 14 days.

### CRISPR/Cas9 gene editing

For CRISPR/Cas9 gene editing, the lentiCRISPR v2-Blast lentiviral vector was modified to contain previously characterized sgRNA sequences ^10^ targeting *CDC42BPA* or *CDC42BPB*. Lentiviral particles were packages and transduced into cells as described above.

### Inducible shRNA knockdown

For shRNA inducible knockdown, the Tet-pLKO-Puro lentiviral vector was modified to contain a shRNA sequences targeting ARHGEF2 (GEF-H1) as described in Supplemental Methods Table 1. Lentiviral particles were packaged and transduced as described above.

### Spheroid formation analysis

OVCAR8 cells were plated at a density of 50,000 per well in 24-well ultralow attachment plates (Corning 3473). BDP9066 was added to cells at the time of plating, and images were taken 24 and 48 hours later using an EVOS digital inverted microscope. Spheroid area was quantitated in ImageJ using the Analyse Particles tool after the colour threshold was set to identify larger clusters of cells.

### Collagen invasion assay

To form spheroids, 500 cells were seeded into 96 well round bottom low attachment plates and allowed to incubate for 48 hours. Collagen I (Gibco) was prepared following the gelling procedure according to manufacturer’s protocol. Spheroids were embedded in 2 mg/mL rat tail collagen I mixture in 6 well plates at 8 spheroids per well. Following polymerization, the collagen matrix was overlaid with 2 mL media containing 2% FBS and, where indicated, drug. The spheroids were imaged immediately following addition of media to record the initial area of the core spheroid and again after 24 h. Invasion was quantified using ImageJ by dividing the area of invasion at 24 h by the initial area of the core spheroid.

### Reverse invasion assay

Complete Matrigel (Avantor) was diluted in PBS (1:1) and 100 µL was pipetted into 6.5 mm diameter Transwell inserts with 8.0 µm pore polycarbonate membranes (Corning). The Matrigel-PBS was left to solidify at 37°C for 30 minutes, then the 24 well plates containing the Transwells were inverted. mCherry-CAAX-expressing OC1 cells were seeded at 6x10^4^ on the membrane at the bottom of the Transwell. Cells were given four hours to adhere, then the plate and Transwells were turned right side up. Cells were washed twice by dipping the Transwells in 1 mL PBS. BDP9066 (3 µM) or the vehicle control DMSO (0.1%) were diluted in supplemented RMPI media (10% FBS, 1 Pen/Strep, 1% Glutamine, 5 µg/mL Insulin) and then added on top of the solidified Matrigel. Cells were left to invade the Matrigel for 5 days. Media and drug were refreshed after 2.5 days. Cells were fixed in 4% para-formaldehyde diluted in PBS. Cells were imaged on a Leica Stellaris 2 confocal microscope at 40x with Type F immersion using diode 638. Optical slices were collected on the Z axis at 10 µm intervals (2 images per Transwell). The mean pixel intensity was measured using Fiji/Image J and averaged from 2 images per replicate. The invert LUTs function was used to invert the images to black on white backgrounds.

### Patient-derived organoid drug screening

Four human high grade serous ovarian cancer patient-derived organoid (PDO) models (OPTO.98, OPTO.129, OPTO.138 or OCAD36.g1) were generated, propagated and screened in the Princess Margaret Cancer Centre Living Biobank core facility (https://pmlivingbiobank.uhnresearch.ca/). PDOs were dissociated to single cells, counted, and seeded onto a thin layer of Matrigel in 384 well plates (2,000 cells per well) 24 h prior to drug treatment ^68^. Organoids were treated with a range of drug concentrations (BDP9066 0.01–50 μM; carboplatin 0.00867-200 μM) or DMSO vehicle for 6 days, and cell viability was assessed using the CellTiter-Glo 3D assay (Promega, USA) according to manufacturer’s instructions. Relative EC_50_ values were calculated from 21-point drug concentrations with four-parameter variable slope nonlinear curve fitting using GraphPad Prism 10.5.0. The human-derived organoid experiments were approved by the Ontario Cancer Research Ethics Board (OCREB) of the Ontario Institute for Cancer Research (OICR) (reference number 0875).

### Protein expression and purification

For bacterial expression of the C3 RhoA inhibitor protein, pGEX 6P1 PTD FLAG C3 was transformed into Rosetta2(DE3)^pLysS^ competent cells (Novagen), and a 1 L culture was grown at 37 °C with 50 µg/mL ampicillin and 34 µg/mL chloramphenicol to an OD_600_ ∼0.8, induced with 0.1 mM IPTG and left to grow for 3 h at 37 °C. Cells were pelleted by centrifugation, resuspended in Tris-buffered saline (TBS) lysis buffer with 2 mM DTT, and lysed by sonication. Lysates were clarified by centrifugation at 10,000 *X* g for 1 h, then incubated with 0.5 mL glutathione-Sepharose 4B resin (Cytiva) overnight at 4 °C with rotation. Beads were washed with 10 column volumes of TBS lysis buffer, then 20 column volumes of Prescission Protease Buffer (50 mM Tris-HCL pH 7.0, 150 mM NaCl, 1 mM EDTA, 1 mM DTT). Beads were incubated overnight with PreScission protease (Cytiva) in Prescission Protease Buffer and the supernatant containing PTD FLAG C3 protein was collected. Protein concentration was determined by using the Bradford Reagent (BioShop).

For expression and purification of Glutathione-S-Transferase (GST), GST-G17A RhoA, and GST-Rhotekin Rho Binding Domain (Rhotekin RBD), BL21(DE3) competent cells (New England Biolabs) were transformed with the appropriate pGEX Construct (GST – pGEX KG ^69^, GST-G17A RhoA ^39^, GST-Rhotekin RBD ^70^). Bacterial cultures were inoculated, grown in 500 mL cultures, and induced as described in García-Mata *et al*., ^39^. Cells were pelleted by centrifugation, resuspended in bacterial lysis buffer (50 mM Tris-HCl, pH 7.5, 150 mM NaCl, 5 mM MgCl_2_, 1% Triton X-100, 1 mM DTT, cOmplete EDTA-free protease inhibitor tablet (Roche)), and lysed by sonication. Lysates were clarified by centrifugation at 20,000 X g for 15 minutes, then incubated with 0.5 mL of glutathione-Sepharose ® 4B resin for 1 h at 4 °C with rotation. Beads were washed with 40 bed volumes of bacterial lysis buffer, then 40 bed volumes of wash buffer (lysis buffer without protease inhibitors). Finally, the beads were resuspended in 1 mL of wash buffer supplemented with 33% glycerol. The bead slurry with bound GST/GST-fusion protein was then aliquoted and stored at -80°C.

GST and GST-fusion proteins were quantified using gel quantitation. Bovine serum albumin (BSA) and samples of purified protein were prepared as SDS-PAGE samples and increasing quantities of each sample were loaded onto a Bis-Tris gradient (4-12%), 1.0 mm, mini protein gel (ThermoFisher). Proteins were visualized by staining with SeeBand Forte solution (FroggaBio) overnight and scanning with the LiCor Odessey Clx according to the manufacturer’s instructions. ImageStudio v6.0 software was used to calculate the densitometry of protein bands.

### Western blotting

Cells were washed with phosphate-buffered saline (PBS) or TBS for phosphorylation-specific work and harvested by cell scraping in the presence of a generic mammalian lysis buffer (TBS pH 7.6, 1% Triton X-100 (v/v), 10 mM EDTA, and cOmplete EDTA-free protease inhibitors). When using phosphorylation-specific antibodies, this buffer was supplemented with phosphatase inhibitors (20 mM NaF, 20 mM β-glycophosphate, and 0.2 μM Na_3_VO_4_). Cells were incubated at 4 °C in lysis buffer with rotation, then clarified by centrifugation at 21,300 X g for 10 minutes at 4 °C.

Alternatively, cells were washed and harvested by cell scraping in the presence of mammalian lysis buffer (1% Sodium lauryl sulfate (SDS), 20 mM Tris-HCl pH 7.5, cOmplete EDTA-free protease inhibitors), then boiled for 5 minutes at 95 °C and lysed by sonication. Lysates were clarified by centrifugation at 21,300 X g for 10 minutes at room temperature.

Protein concentrations were determined using the Pierce BCA Protein Assay Kit (ThermoFisher) according to the manufacturer’s instructions. Normalized lysates were prepared as western blot samples by diluting them by 25% with 4X Bolt LDS Sample Buffer (Invitrogen) and 10% with β-mercaptoethanol (BME – Sigma Aldrich). Samples were boiled at 95 °C for 5 minutes. Proteins were resolved by SDS-PAGE on Bis-Tris 4-12% gradient protein gels, then transferred onto 0.45 μM nitrocellulose membranes (Bio-Rad). Membranes were blocked with a solution of 3% BSA in TBS with 0.1% TWEEN-20 (BioShop) (TBS-T) for 1 hour, then incubated with primary antibodies (described in Supplemental Methods Table 1) in the blocking solution, overnight with gentle agitation. Membranes were washed 3 times with TBS-T then incubated with secondary antibodies (Supplemental Methods Table 1) at room temperature for 1 h. Membranes were washed with TBS-T and dH_2_O, then visualized on the Licor Odessey CLx instrument according to the manufacturer’s instructions.

### Active protein pulldowns

Cells were cultured and manipulated as indicated, then lysed as described previously. For active RhoA pulldowns, cells were lysed in a GTPase stable lysis buffer (TBS pH 7.6, 1% Triton X-100 (v/v), 10 mM EGTA (BioShop), 10 mM MgCl2, and protease inhibitors). In all other cases, cells were lysed in generic lysis buffer, with the addition of phosphatase inhibitors for phosphorylation-specific assays. Lysates were collected and immediately distributed to 25 μg of GST or GST-fusion proteins on Glutathione Sepharose beads (prepared previously, as described). Active proteins were selectively precipitated using their GST-fused binding partners (*e.g*. active RhoA : GST-Rhotekin RBD; active GEF-H1 : GST-G17A RhoA), with GST alone used as a negative control. Lysates and beads were incubated at 4 °C for 1 h with rotation. Beads were washed 3-5 times with lysis buffer, then resuspended in 50 μL of lysis buffer, prepared as an SDS-PAGE sample, and analysed via western blotting.

### Co-immunoprecipitations

Cells were cultured and manipulated as desired, then lysed as described previously in generic lysis buffer. 300 μL of lysate was incubated overnight with 1.25 µg of the IP antibody or isotype control (Supplemental Methods Table 1) at 4 °C with rotation. The antibody-lysate complexes were then incubated for 4 h at 4 °C with 18.75 μL of Protein A/G Agarose beads (Calbiochem, IP05), then the beads were pelleted by centrifugation, washed 3 times with the lysis buffer, prepared as an SDS-PAGE sample, and analysed by western blotting.

For the α-tubulin immunoprecipitation, additional components were added to the IP lysis buffer (100 mM PIPES pH 6.9, 0.1% Triton-X100, 0.1% IGEPAL-CA630 NP40 alternative (Sigma Aldrich), 0.1% TWEEN-20, 30% Glycerol, 1mM EGTA, 5 mM MgCl2, 0.1% BME, protease inhibitors) to facilitate protein stability. To prevent nonspecific interactions, 2 µg of the desired or control IP antibody was first incubated with Protein A/G Agarose beads at 4 °C overnight with rotation. Cells were lysed and lysates were pre-cleared by incubation with 20 μL of Protein A/G beads without antibodies, for 30 minutes at 4 °C with rotation. Pre-cleared lysates were transferred to the Protein A/G bead-antibody complexes and left to incubate for 1 h at 4 °C with rotation. The beads were washed 5 times with IP lysis buffer, then prepared as an SDS-PAGE sample and analysed by western blotting.

### In vitro kinase assay and mass spectrometry

One μg of recombinant MRCKα kinase domain (1-473 aa) (Baculovirus, Active Motif, 81396) or full length purified CMV-FLAG-*CDC42BPA* and 2.5 μg purified GST-GEF-H1 substrate were added to a buffer containing 20 mM Tris-HCl pH 7.5, 0.01% Tween20, 1 mM MgCl_2_, and 0.5 mM ATP in a final volume of 40 μl. The reaction was incubated at 30°C at 1300 rpm for 1 h in an incubator shaker. The reaction was boiled for 5 min at 100°C to stop the reaction. After cooling, one volume (40 μL) of 50 mM ammonium bicarbonate was added along with 25 ng trypsin. The reaction was digested overnight at 37°C with shaking. The following day, digested peptides were isolated using C-18 stage tips. Proteolytic peptides were resuspended in 0.1% formic acid and separated with a Thermo Scientific RSLC nano Ultimate 3000 LC on a Thermo Scientific Easy-Spray C-18 PepMap 75 µm x 50 cm C-18 2 μm column. A 75 min gradient of 2-25% (30 min) then 25%-90% (5 min) acetonitrile with 0.1% formic acid was run at 300 nL/min at 50°C. Eluted peptides were analyzed with a Thermo Scientific Orbitrap Exploris 480 mass spectrometer utilizing a top 15 methodology in which the 15 most intense peptide precursor ions were subjected to fragmentation. The normalized AGC target for MS1 was set to 300% with a max injection time of 25 ms, the normalized AGC target for MS2 ions was set to 100% with a max injection time of 40 ms, and the dynamic exclusion was set to 42 s.

### GST-protein purification and TEV cleavage for soluble protein

GST-*ARHGEF2* (EX-Z7233-B03-9181) was purchased from GeneCopoeia. Plasmid DNA was transformed in BL21 (DE3) Competent *E. Coli* (New England Biolabs-C2527H) for use of T7-based bacterial expression. Carbenicillin cultures (100 mL LB + 100 μg/mL) were inoculated in a 25°C shaker (250 RPM) until O.D. = 0.6 followed by IPTG induction overnight at 25°C. Cultures were harvested by centrifugation at 2400 x g for 20 min at 4°C. Supernatant was discarded and pellets lysed in 3 mL of Complete NETN buffer (20 mM 1M Tris pH 8.0, 100 mM NaCl, 1 mM EDTA, 0.5% NP-40, 500 µM PMSF, 1 µg/mL Leupeptin). Lysates were sonicated using a Branson Sonifier 250 at 30% duty cycle, output control level 2, using 15 s on + 45 s off cycles for 30 cycles. GST-tagged proteins were pulled down using 100 μL of GenScript Glutathione MagBeads (Cat. L00895-2), prewashed with complete NETN buffer (50% slurry) and allowed to rotate overnight, incubating at 4°C. Protein loaded beads were washed twice (1 mL) with Complete NETN containing 0.7 M NaCl (High-Salt), followed by 2 low-salt Complete NETN washes. Protein loaded beads were subjected to a buffer exchange for TEV Cleavage 50 mM Tris-HCl (pH 7.5), 0.5 mM EDTA, and 1 mM DTT and incubated with 20 μg of His-tagged Super-TEV overnight rotating in a 250 µL volume. Cleaved protein was collected and the Super-TEV removed by GenScript Ni-Charged MagBead (Cat.L00295) cleanup (50 µL of TEV-Buffer Washed beads - 50% slurry) for 2 h rotating at 4°C and the supernatant collected as the purified soluble protein and analyzed for downstream applications.

### FLAG-tag protein purification

CMV-FLAG-*CDC42BPA* was purchased from GeneCopoeia. Five 150 mm plates of HEK293T cells were transfected at 80% confluency using Lipofectamine 3000 (45 µg of plasmid DNA: 90 µL p3000 reagent: 90 µL Lipofectamine 3000) and allowed to express for 48 h. Each plate of cells was lysed in MIB buffer (750 µL) and collected by scraping. Centrifugation was carried out at 17000 x g for 45 minutes and supernatants were placed into clean microcentrifuge tubes for individual immunoprecipitations. FLAG-tagged proteins were immunoprecipitated by rotating overnight at 4°C using 60 µL (50% slurry of prewashed Pierce™ Anti-DYKDDDDK Magnetic Agarose (A36797) in MIB buffer). Supernatants were collected and beads pooled to carry out washes in 1.5 mL microcentrifuge tubes. Fifteen washes (1 mL) were performed using low-salt MIB buffer. FLAG-protein loaded beads were eluted twice using 200 µg/mL of 3x FLAG peptide (Sigma F4799) in a 150 µL volume of complete MIB buffer (300 µL total). FLAG peptide was removed using Pierce™ Protein Concentrator, 50K MWCO columns (ThermoFisher 88504) until 100 µL remained. Glycerol and DTT were added to the preparation for protein stability at final concentrations of 25% and 1 mM respectively. Final concentrations of the purified proteins were determined using BCA.

### AlphaFold2 modeling of GEF-H1 phosphorylation

Full-length AlphaFold2 prediction of ARHGEF2 (GEF-H1) AF-Q92974-F1-model_v4 was downloaded from UniProt and rendered in Pymol 3.0.5, colored by pLDDT scoring utilized by AlphaFold2. Sites of phosphorylation (S121, S122, S174, and S886) are circled, colored in black, and the residue numbers annotated as determined by MRCKα *in vitro* kinase assays using purified MRCKα full-length and kinase domain.

### FLAG immunoprecipitation and mass spectrometry

Cells were transfected using Lipofectamine 3000 48 hours prior to harvesting according to manufacturer’s instructions. Cells were harvested in a buffer containing 25 mM TrisCl (pH 7.5), 1% NP-40, 150 mM NaCl, 1 mM EDTA, 5% glycerol, 1X protease inhibitor cocktail (Roche), and 1x each of phosphatase inhibitor cocktails 2 and 3 (Sigma). Lysates were incubated for 30 minutes using over-end rotation, then centrifuged at 21,000 rpm for 15 minutes at 4°C. Anti-DYKDDDDK magnetic beads (Pierce) were added to the lysates at a ratio of 10 μL beads per 1 mg of lysate and incubated for 2 hours using over-end rotation. Beads were washed 4x with lysis buffer, then boiled in Elution Buffer (0.5% SDS, 1% BME, 0.1 M Tris-HCl pH 6.8). Eluted proteins were reduced by incubation with 5 mM DTT at 65°C for 25 minutes and alkylated with 20 mM iodoacetamide at room temperature for 30 minutes in the dark. Samples were concentrated to 100 µL with Millipore 10kD cutoff spin concentrators. Detergent was removed by chloroform/methanol extraction and protein pellets were digested trypsin overnight at 37°. Peptides were cleaned with C18 stage tips (Thermo), then resuspended in 0.1% formic acid and separated with a Thermo Scientific RSLCnano Ultimate 3000 LC on a Thermo Scientific Easy-Spray C-18 PepMap 75 µm x 50 cm C-18 2 μm column. A 240 min gradient of 2-25% (30 min) then 25%-90% (5 min) acetonitrile with 0.1% formic acid was run at 300 nL/min at 50°C. Eluted peptides were analyzed by a Thermo Scientific Orbitrap Exploris 480 mass spectrometer utilizing a top 20 methodology in which the 20 most intense peptide precursor ions were subjected to fragmentation. The normalized AGC target for MS1 was set to 300% with a max injection time of 25 ms, the normalized AGC target for MS2 ions was set to 100% with a max injection time of 22 ms, and the dynamic exclusion was set to 60 s.

### Live cell confocal videomicroscopy and analysis

OVCAR8 cells stably expressing GFP-RhoA and Lifeact-TagRFPt for live cell imaging were passaged 24-48 hours post-transfection cells with 0.05% Trypsin/EDTA, suspended in phenol red free RPMI 1640 Medium (supplemented with 10% FBS) and seeded at approximately 50,000 to 100,000 cells per well in 6-well dish on 18 mm diameter, #1 -thickness circular glass coverslips (Electron Microscopy Sciences). Cells were cultured to target 20-30% confluency 48-72 hours post-seeding. Thirty minutes to one hour before imaging, coverslips were transferred to a preheated magnetic imaging chamber containing 800 µl phenol red-free RPMI supplemented with 10% FBS and mounted in a temperature- and CO₂-controlled incubation chamber (37 °C, 5% CO₂). Time-lapse imaging was performed for 1–2 h on a Leica DMi8 inverted microscope equipped with an Andor Diskovery multimodal spinning-disk system, an Andor iXON 897 EMCCD camera, and a Leica 63X oil-immersion objective (NA 1.518), using Quorum Wave FX powered by MetaMorph software. Z-stacks (7 µm total depth, 0.5 µm step size) were acquired every 2 min. GFP and RFP signals were excited at 488 nm and 561 nm, respectively, with laser power adjusted (5–8%) to minimize phototoxicity. Fields of view were selected to include two or more cells in close proximity and likely to establish cell–cell contacts during acquisition. Upon observation of a newly formed cell–cell junction (typically 10–20 min after imaging start), cells were treated with either DMSO (vehicle) or BDP9066 (3 µM final concentration). Drug addition was performed during a 2-min imaging interval by gently adding 200 µl of pre-warmed medium containing DMSO or BDP9066, bringing the total volume to 1 ml. Imaging was continued for at least 60 min after treatment.

Image processing and analysis were performed using ImageJ/Fiji. Confocal z-stacks were converted to single-plane images using maximum-intensity projection, and each fluorescent channel was processed independently using the optimal z-section for that marker. Nuclear displacement was quantified using the eGFP-RhoA channel, which is excluded from the nucleus. Nuclear position was estimated by identifying the z-plane where nuclear boundaries were most clearly defined, and nuclear outlines were manually traced to determine centroid positions at multiple time points before and after drug addition. Inter-nuclear distances between neighboring cells were calculated for each time point and normalized by subtracting the distance measured in the final frame prior to treatment.

### Statistical analysis

GraphPad Prism 10.6.1 was used for all statistical analysis, with details provided in figure legends for each test.

## Author contributions

A.M.K. performed mass spectrometry, analyzed proteomics datasets and submission to PRIDE, Western blotting, immunoprecipitations, RNAi knockdown and spheroid invasion assays. J.S.W. performed protein purification of GEF-H1 and Flag-MRCKα, and GEF-H1 phosphorylation site modeling. I.K.M.K. performed immunoblots, spheroid cell viability and spheroid formation assays. A.R. performed RhoA and GEF-H1 activity assays, tubulin immunopreciptations, and measured GEF-H1 phosphorylation by Western blotting. A.C. performed reverse invasion assays, MRCK/GEF-H1 immunopreciptations, GEF-H1 domain mapping, ERM activation. A.M.S.C. established LifeAct TagRFP-T and GFP-RhoA expressing OVCAR8 cells and performed live-cell imaging. D.O. established MRCK CRISPR/Cas9 knockout OVCAR8 cells and determined the RhoA-dependence for YAP and ERM phosphorylation. R.G. made various DNA constructs. N.R. performed patient-derived organoid viability assays. J.S.D. and M.F.O contributed to conceptualization, project administration, formal analysis, visualization, supervision, funding acquisition and writing, review and editing of the manuscript.

## Acknowledgements

Thanks to Robert Rottapel (Princess Margaret Cancer Centre, Toronto ON, Canada) for the gift of GEF-H1 deletion mutant plasmids ^38^. Thanks to Denise Connolly (Cancer Signaling and Microenvironment, Fox Chase Cancer Center, Philadelphia, PA, USA) for providing the OC-1 cell line. lentiCRISPR v2-Blast was a gift from Mohan Babu (Addgene plasmid # 83480; http://n2t.net/addgene:83480; RRID:Addgene_83480). tetO-FUW-eGFP-RHOA-Q63L was a gift from Andrew Putnam (Addgene plasmid # 73081; http://n2t.net/addgene:73081; RRID:Addgene_73081) ^71^. pLenti-mCherry-CAAX PuroR was a gift from Carlos Carmona-Fontaine (Addgene plasmid # 166228; http://n2t.net/addgene:166228; RRID:Addgene_166228) ^72^. pCMV-VSV-G was a gift from Bob Weinberg (Addgene plasmid # 8454; http://n2t.net/addgene:8454; RRID:Addgene_8454) ^73^. psPAX2 was a gift from Didier Trono (Addgene plasmid # 12260; http://n2t.net/addgene:12260; RRID:Addgene_12260). pGEX-4T1-RhoA G17A was a gift from Rafael Garcia-Mata (Addgene plasmid # 69357; http://n2t.net/addgene:69357; RRID:Addgene_69357) ^39^. GST-Rhotekin RBD was a gift from Martin Schwartz (Addgene plasmid # 15247; http://n2t.net/addgene:15247; RRID:Addgene_15247) ^70^. The Tet-pLKO-puro lentiviral vector was gift from Dmitri Wiederschain (Addgene plasmid # 21915, https://www.addgene.org/21915/; RRID:Addgene_21915) ^74^.

