## Supplemental Figures for "MRCKα represses GEF-H1 mediated RhoA activation to promote ovarian cancer spheroid growth and invasion"


| Supplemental Figures | Page |
|---|---|
| **Figure S1. Identification of MRCKα interacting proteins in HGSOC cells.** | **2** |
| **Figure S2. MRCKα interacts with the GEF-H1 N-terminus.** | **4** |
| **Figure S3. MRCK deletion or inhibition increases active GEF-H1 available for binding to nucleotide-free RhoA.** | **6** |
| **Figure S4. MRCK inhibition activates RhoA.** | **8** |
| **Figure S5. FGD1 induction results in CDC42 activation and MRCK inhibitor-sensitive reduction in RhoA activity.** | **10** |
| **Figure S6. RhoA activation induced by MRCK deletion or inhibition is GEF-H1 dependent.** | **12** |
| **Figure S7. Mass spectrometry characterization of GEF-H1 phosphorylation by MRCKα.** | **14** |
| **Figure S8. MRCK phosphorylation of GEF-H1 on Serine 174.** | **16** |
| **Figure S9. MRCK inhibition reduces GEF-H1 association with α-Tubulin.** | **18** |
| **Figure S10. Decreased patient derived organoid viability induced by BDP9066 was not observed for the less potent MRCK inhibitor BDP9069 enantiomer.** | **20** |
| **Video 1. Time-lapse microscopy of non-targeting control (NTC) siRNA transfected OVCAR8 cells treated with BDP9066.** | **22** |
| **Video 2. Time-lapse microscopy of GEF-H1-targted siRNA transfected OVCAR8 cells treated with BDP9066.** | **22** |
| **Video 3. Time-lapse microscopy of NTC transfected OVCAR8 cells treated with DMSO.** | **22** |

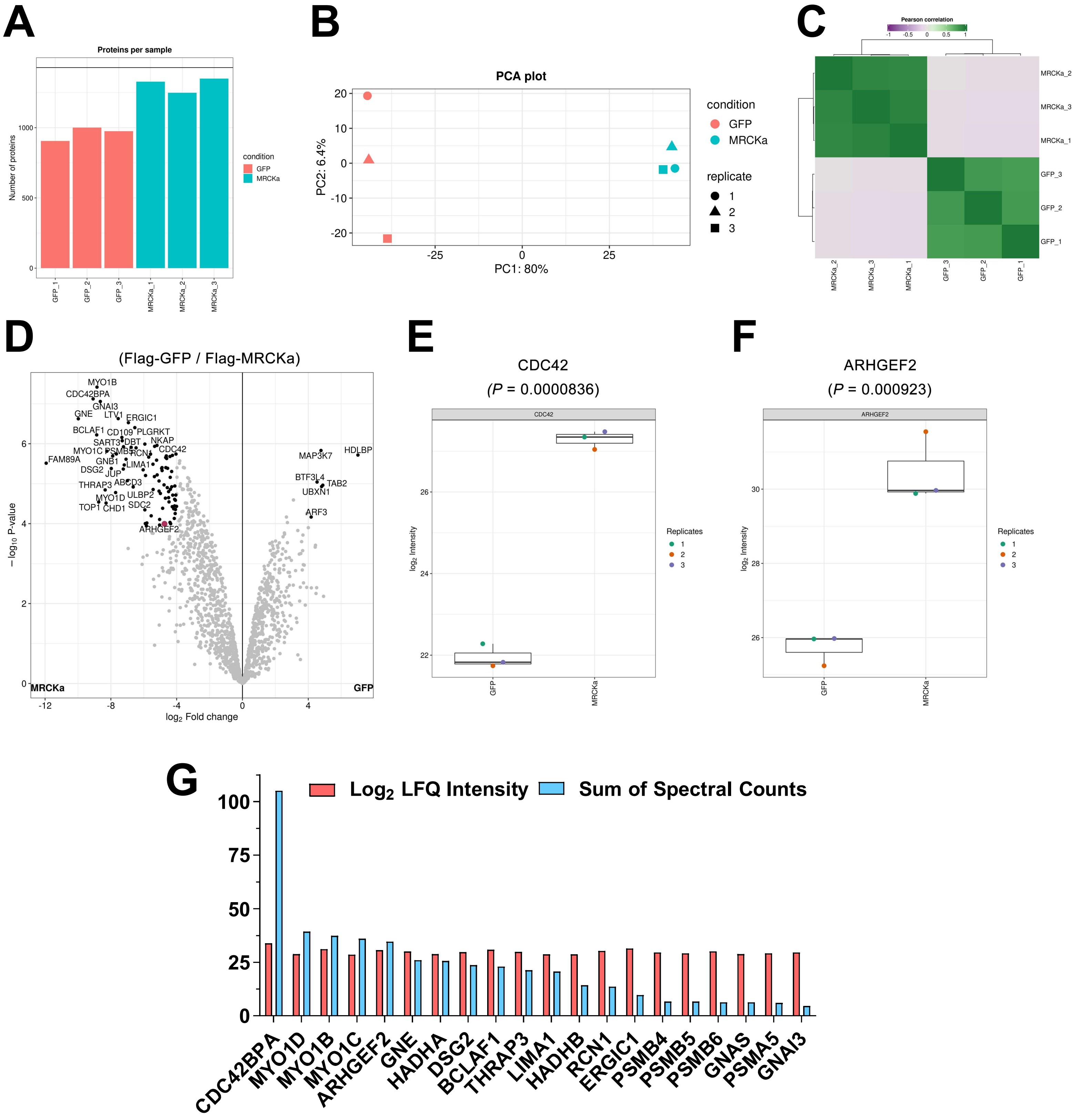
A
Proteins per sample
Number of proteins
condition
GFP
MRCKa
B
PCA plot
PC1: 80%
PC2: 6.4%
condition
GFP
MRCKa
replicate
C
Pearson correlation
D
(Flag-GFP / Flag-MRCKa)
log2 Fold change
− log10 P-value
MRCKa
GFP
E
CDC42
(P = 0.0000836)
F
ARHGEF2
(P = 0.000923)
Replicates
G
Log2 LFQ Intensity
Sum of Spectral Counts
CDC42BPA
MYO1D
MYO1B
MYO1C
ARHGEF2
GNE
HADHA
DSG2
BCLAF1
THRAP3
LIMA1
HADHB
RCN1
ERGIC1
PSMB4
PSMB5
PSMB6
GNAS
PSMA5
GNAI3

**Figure S1**

**Figure S1. Identification of MRCKα interacting proteins in HGSOC cells. A**. Total number of proteins identified in each FLAG-MRCKα or FLAG-GFP replicate immunoprecipitation from OC-1 cells. **B**. PCA plot analysis comparing proteins enriched by FLAG-MRCKα or FLAG-GFP replicate immunoprecipitations from OC-1 cells. **C**. Pearson correlation of proteins enriched by FLAG-MRCKα or FLAG-GFP replicate immunoprecipitations from OC-1 cells. **D**. Volcano plot depicts protein abundance comparing proteins enriched by FLAG-MRCKα or FLAG-GFP immunoprecipitation from OC-1 cells. CDC42BPA = MRCKα. Raw LFQ intensity of **E.** CDC42 or **F**. ARHGEF2 (GEF-H1) enriched in replicate immunoprecipitations from OC-1 cells. **G**. Bar graph depicts the top 20 most abundant proteins enriched by FLAG-MRCKα relative to FLAG-GFP immunoprecipitation from OC-1 cells, as determined by LFQ intensity or the sum of spectral counts.

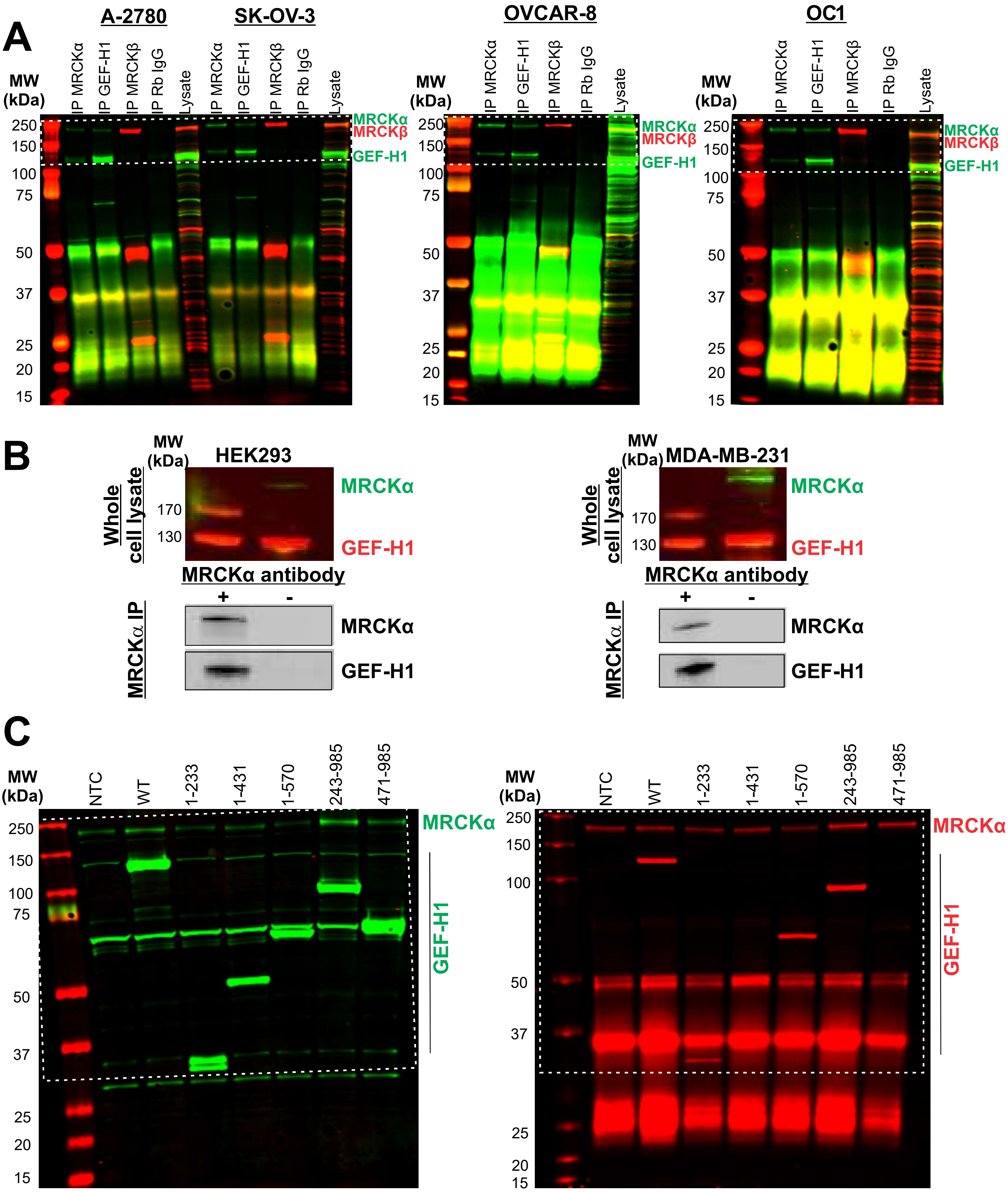
A
A-2780
SK-OV-3
OVCAR-8
OC1
MW (kDa)
IP MRCKα
IP GEF-H1
IP MRCKβ
IP Rb IgG
Lysate
MRCKα
MRCKβ
GEF-H1
250
150
100
75
50
37
25
20
15
B
HEK293
MDA-MB-231
Whole cell lysate
170
130
MRCKα antibody
+
-
MRCKα IP
C
NTC
WT
1-233
1-431
1-570
243-985
471-985

Figure S2

**Figure S2. MRCKα interacts with the GEF-H1 N-terminus. A.** White rectangles with dotted borders indicate cropped regions used in Figure 3A. **B.** Cell lysates from HEK293 human embryonic kidney (left panels) or MDA-MB-231 human breast cancer cells (right panels) were western blotted with antibodies against MRCKα (green) and GEF-H1 (red). Lysates were incubated with (+) or without (-) antibodies against MRCKα, immunoprecipitated proteins were western blotted with antibodies against MRCKα and GEF-H1. **C.** White rectangles with dotted borders indicate cropped regions used in Figure 3B.

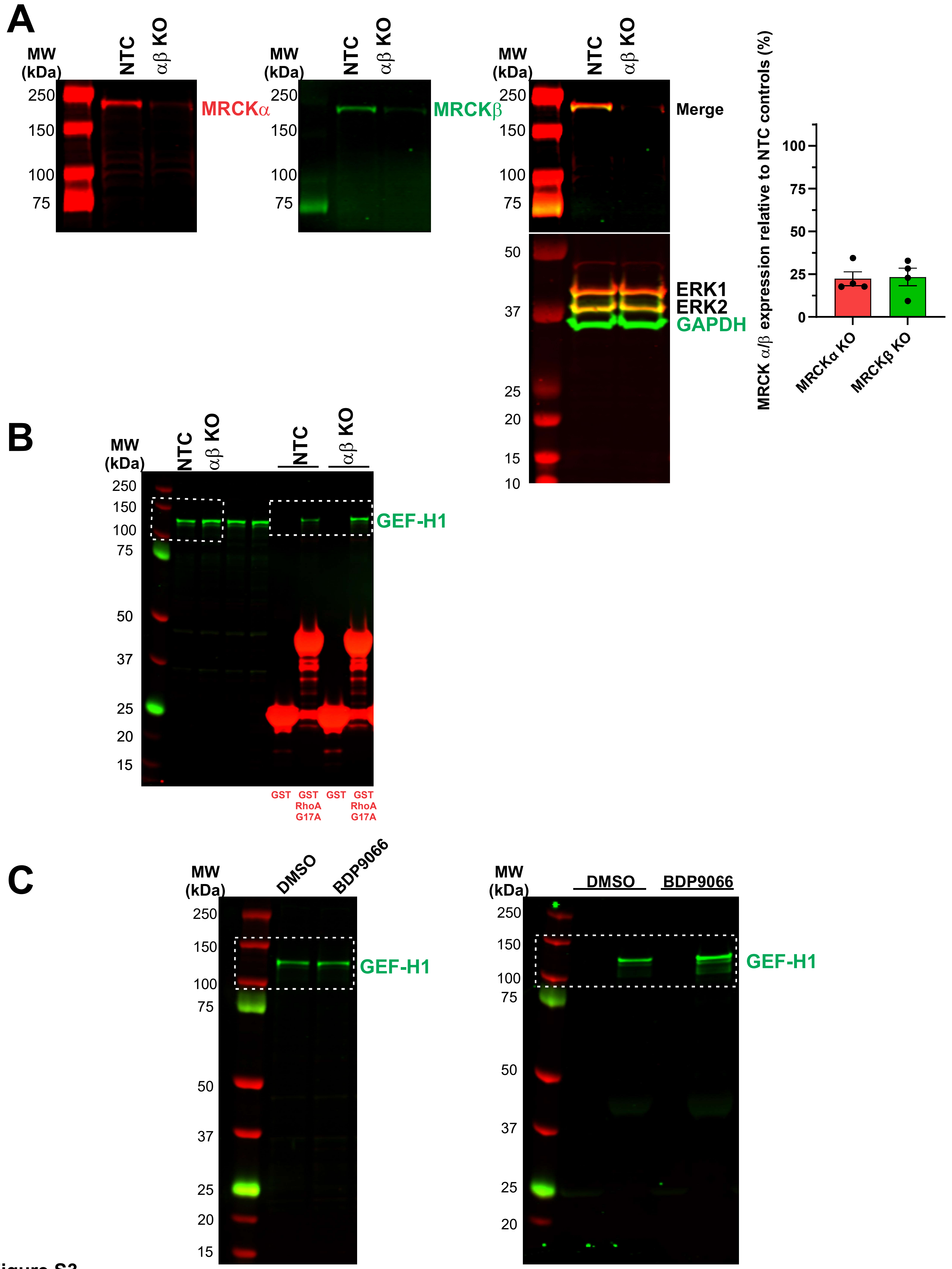
A
MW (kDa)
NTC
αβ KO
250
150
100
75
MRCKα
MRCKβ
Merge
50
37
25
20
15
10
ERK1
ERK2
GAPDH
MRCK α/β expression relative to NTC controls (%)
100
75
50
25
0
MRCKα KO
MRCKβ KO
B
GEF-H1
GST
GST RhoA G17A
C
DMSO
BDP9066

Figure S3

**Figure S3. MRCK deletion or inhibition increases active GEF-H1 available for binding to nucleotide-free RhoA. A.** Non-targeted control (NTC) or double knockout αβ KO OVCAR8 cells were quantitatively western blotted with antibodies specific for MRCKα (red) and MRCKβ (green) as indicated. Antibodies against GAPDH and ERK1/ERK2 were used as loading controls. The ratios of MRCKα or MRCKβ were normalized to NTC cells for each independent replicate. Means ± SEM, N = 4. White rectangles with dotted borders indicate cropped regions used in **B.** Figure 4A. **C.** Figure 4B.

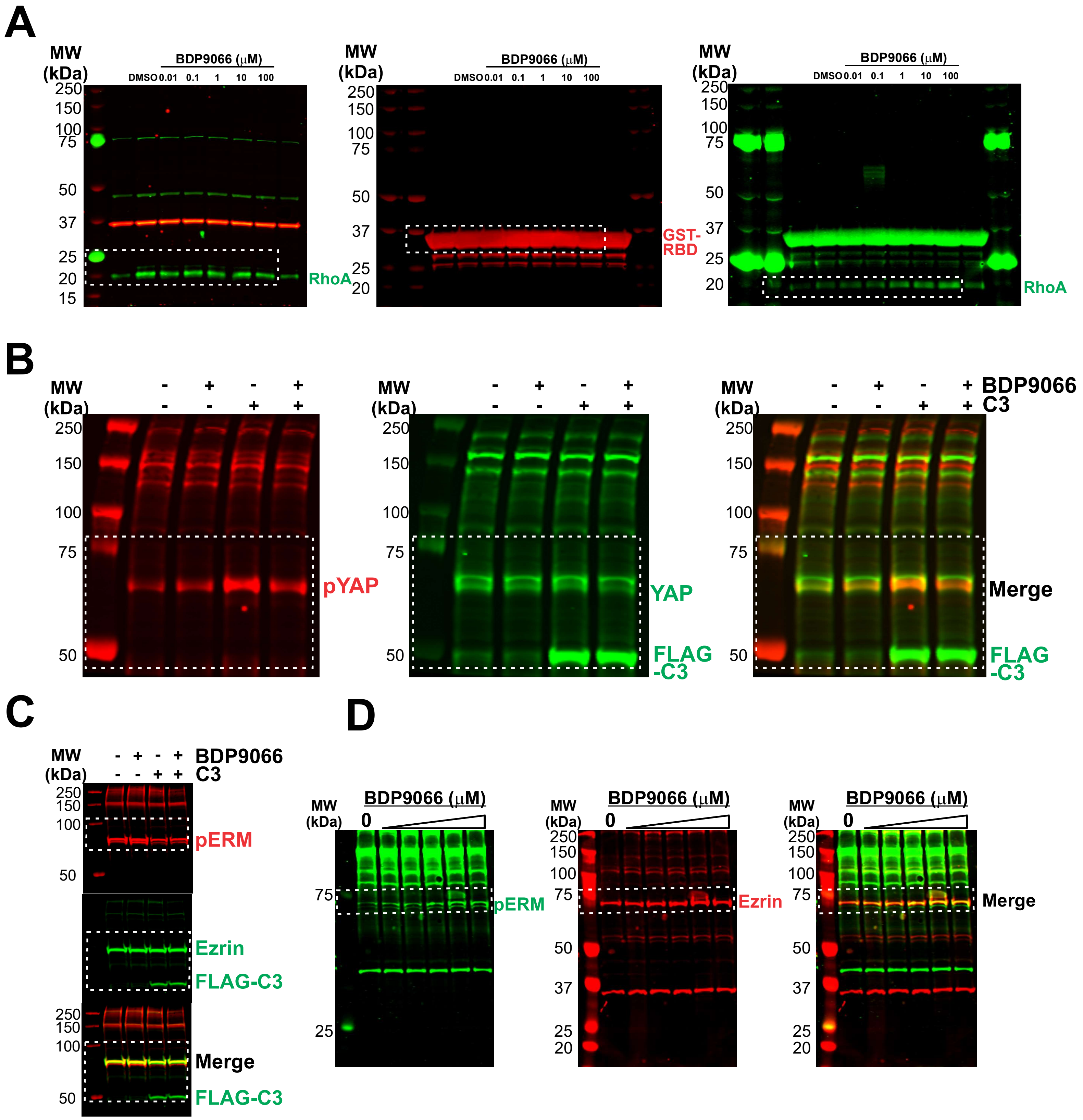
A
MW (kDa)
BDP9066 (μM)
DMSO 0.01 0.1 1 10 100
250 150 100 75 50 37 25 20 15
RhoA
GST-RBD
B
BDP9066
C3
pYAP
YAP
FLAG -C3
Merge
C
pERM
Ezrin
FLAG-C3
D
0


Figure S4

**Figure S4. MRCK inhibition activates RhoA. A.** White rectangles with dotted borders indicate cropped regions used in Figure 5A. **B.** Figure 5B, **C.** Figure 5C, **D.** Figure 5D.

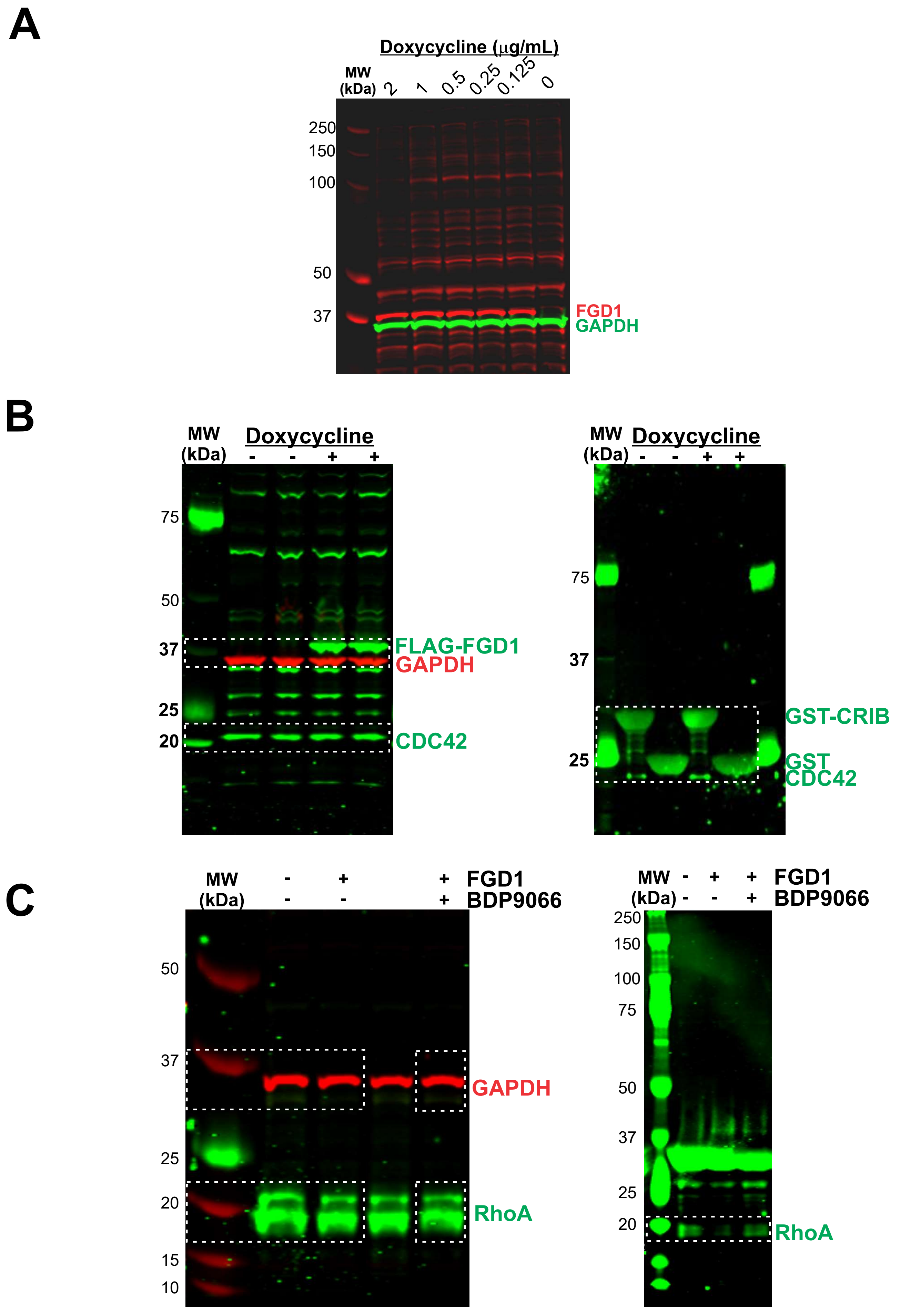

A
Doxycycline (μg/mL)
MW (kDa)
2
1
0.5
0.25
0.125
0
250
150
100
50
37
FGD1
GAPDH
B
MW (kDa)
Doxycycline
- - + +
75
50
37
25
20
FLAG-FGD1
GAPDH
CDC42
MW (kDa)
Doxycycline
- - + +
75
37
25
GST-CRIB
GST
CDC42
C
MW (kDa)
- + + FGD1
- - + BDP9066
50
37
25
20
15
10
GAPDH
RhoA
MW (kDa)
- + + FGD1
- - + BDP9066
250
150
100
75
50
37
25
20
RhoA


Figure S5

**Figure S5. FGD1 induction results in CDC42 activation and MRCK inhibitor-sensitive reduction in RhoA activity. A.** OVCAR8 cells transfected with doxycycline-inducible FLAG-tagged FGD1 were treated with indicated doxycycline concentrations for 48 h. White rectangles with dotted borders indicate cropped regions used in **B.** Figure 5E, **C.** Figure 5F.

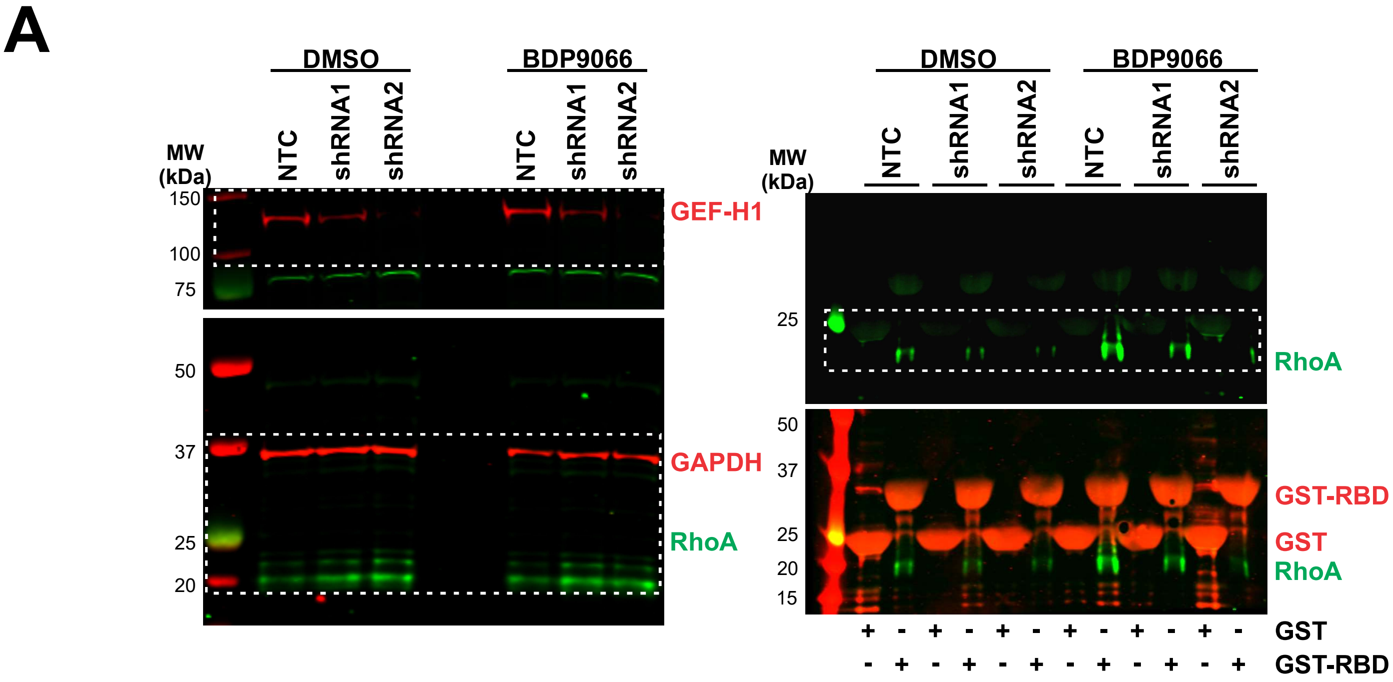
A
DMSO
BDP9066
NTC
shRNA1
shRNA2
MW (kDa)
150
100
75
50
37
25
20
15
GEF-H1
GAPDH
RhoA
GST-RBD
GST
+ - + - + - + - + - + -
- + - + - + - + - + - +

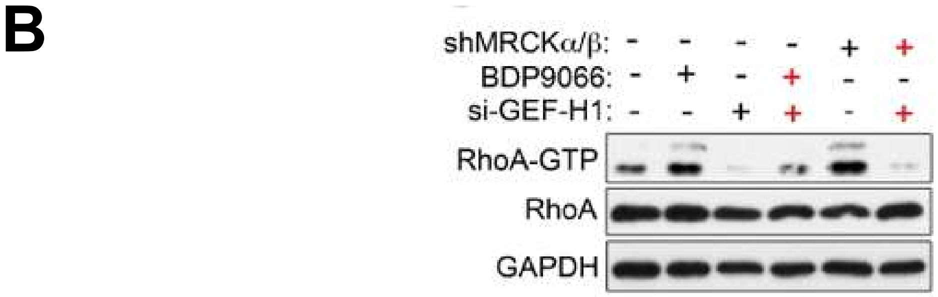
B
shMRCKα/β: - - - - + +
BDP9066: - + - + - -
si-GEF-H1: - - + + - +
RhoA-GTP
RhoA
GAPDH

Figure S6

**Figure S6. RhoA activation induced by MRCK deletion or inhibition is GEF-H1 dependent.** White rectangles with dotted borders indicate cropped regions used in **A.** Figure 6A. **B.** Figure 6B. **C.** OC-1 cells in which MRCKα and MRCKβ were knocked down with shRNA as indicated, or were treated with BDP9066, were either left untransfected or transfected with siRNA targeting GEF-H1. GST-RBD was used to affinity purify active RhoA-GTP, which was western blotted with anti-RhoA antibody. In addition, total RhoA and GAPDH were western blotted.

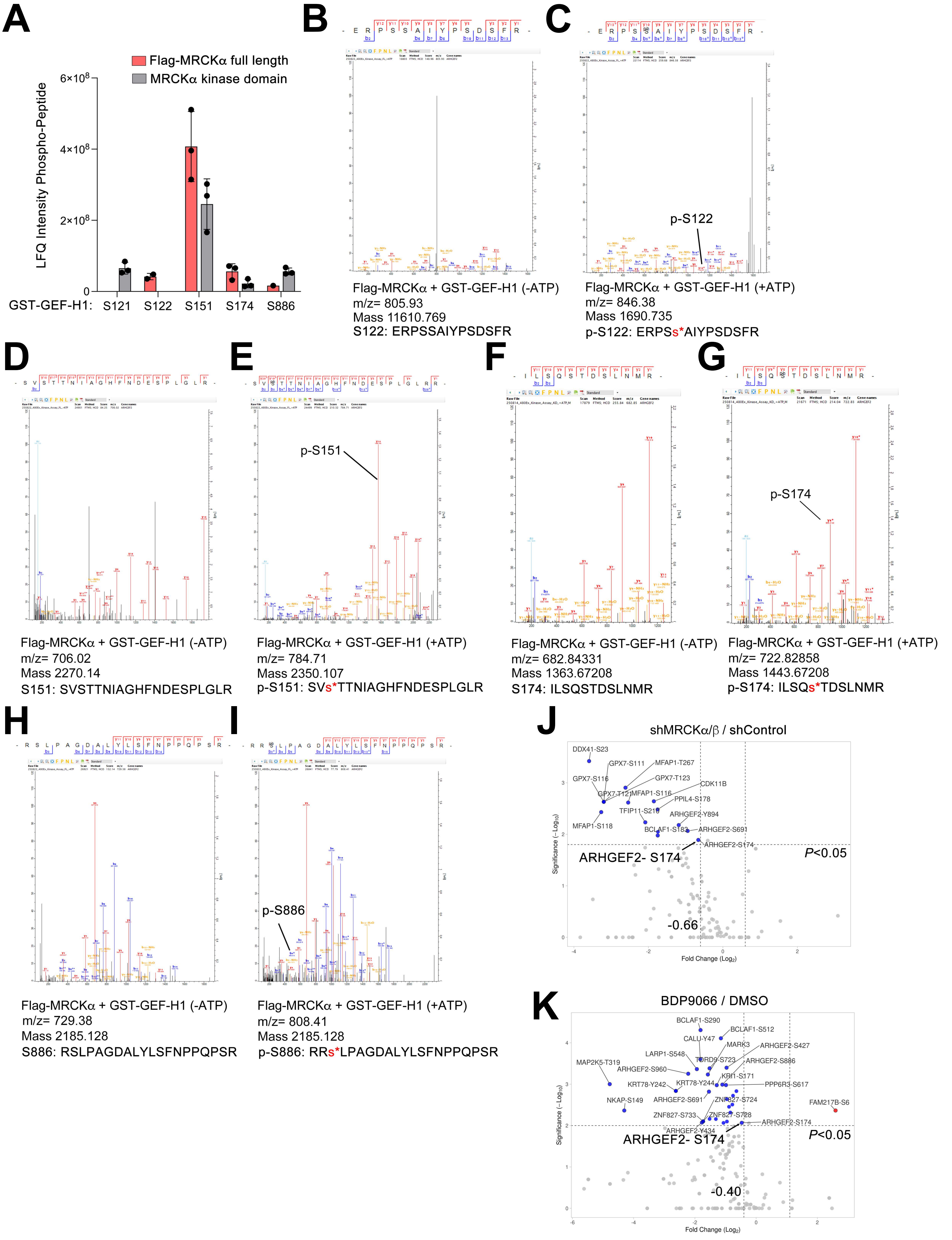
A
Flag-MRCKα full length
MRCKα kinase domain
LFQ Intensity Phospho-Peptide
6×10^8
4×10^8
2×10^8
0
GST-GEF-H1: S121 S122 S151 S174 S886
B
Flag-MRCKα + GST-GEF-H1 (-ATP)
m/z= 805.93
Mass 11610.769
S122: ERPSSAIYPSDSFR
C
p-S122
Flag-MRCKα + GST-GEF-H1 (+ATP)
m/z= 846.38
Mass 1690.735
p-S122: ERPSs*AIYPSDSFR
D
Flag-MRCKα + GST-GEF-H1 (-ATP)
m/z= 706.02
Mass 2270.14
S151: SVSTTNIAGHFNDESPLGLR
E
p-S151
Flag-MRCKα + GST-GEF-H1 (+ATP)
m/z= 784.71
Mass 2350.107
p-S151: SVs*TTNIAGHFNDESPLGLR
F
Flag-MRCKα + GST-GEF-H1 (-ATP)
m/z= 682.84331
Mass 1363.67208
S174: ILSQSTDSLNMR
G
p-S174
Flag-MRCKα + GST-GEF-H1 (+ATP)
m/z= 722.82858
Mass 1443.67208
p-S174: ILSQs*TDSLNMR
H
Flag-MRCKα + GST-GEF-H1 (-ATP)
m/z= 729.38
Mass 2185.128
S886: RSLPAGDALYLSFNPPQPSR
I
p-S886
Flag-MRCKα + GST-GEF-H1 (+ATP)
m/z= 808.41
Mass 2185.128
p-S886: RRs*LPAGDALYLSFNPPQPSR
J
shMRCKα/β / shControl
DDX41-S23
GPX7-S111
MFAP1-T267
GPX7-S116
GPX7-T123
CDK11B
GPX7-T121
MFAP1-S116
PPIL4-S178
TFIP11-S210
ARHGEF2-Y894
MFAP1-S118
BCLAF1-S183
ARHGEF2-S691
ARHGEF2-S174
ARHGEF2- S174
P<0.05
-0.66
Significance (−Log10)
Fold Change (Log2)
K
BDP9066 / DMSO
BCLAF1-S290
BCLAF1-S512
CALU-Y47
MARK3
ARHGEF2-S427
LARP1-S548
TORD9-S723
ARHGEF2-S886
MAP2K5-T319
ARHGEF2-S960
KRI1-S171
KRT78-Y242
KRT78-Y244
PPP6R3-S617
NKAP-S149
ARHGEF2-S691
ZNF827-S724
FAM217B-S6
ZNF827-S733
ZNF827-S728
ARHGEF2-S174
ARHGEF2-Y434
ARHGEF2- S174
P<0.05
-0.40
Significance (−Log10)
Fold Change (Log2)

**Figure S7**

**Figure S7. Mass spectrometry characterization of GEF-H1 phosphorylation by MRCKα. A**. Bar graph depicts LFQ intensities of GST-GEF-H1 peptides phosphorylated by Flag-MRCKα or MRCKα kinase domain (1-473 aa) using *in vitro* kinase assays from three independent replicate experiments. Mean ± SD. **B-I**. Raw MS/MS spectra of non-phosphorylated or phosphorylated GEF-H1 tryptic peptides from *in vitro* kinase assays with Flag-MRCKα and GST-GEF-H1 in the presence or absence of ATP. Volcano plots of GEF-H1 phosphorylated peptides with increased or reduced abundance following **J.** MRCKα/β knockdown by shRNAs or **K**. 5 μM BDP9066-treatment for 24 hours. Differences in $\log_2$ LFQ intensities amongst control or MRCKμ/β knockdown or BDP9066-treatment were determined by paired t test with adjusted p-value cutoff of 0.05 using Perseus Software. Volcano plots were generated using VolcaNoseR [77].

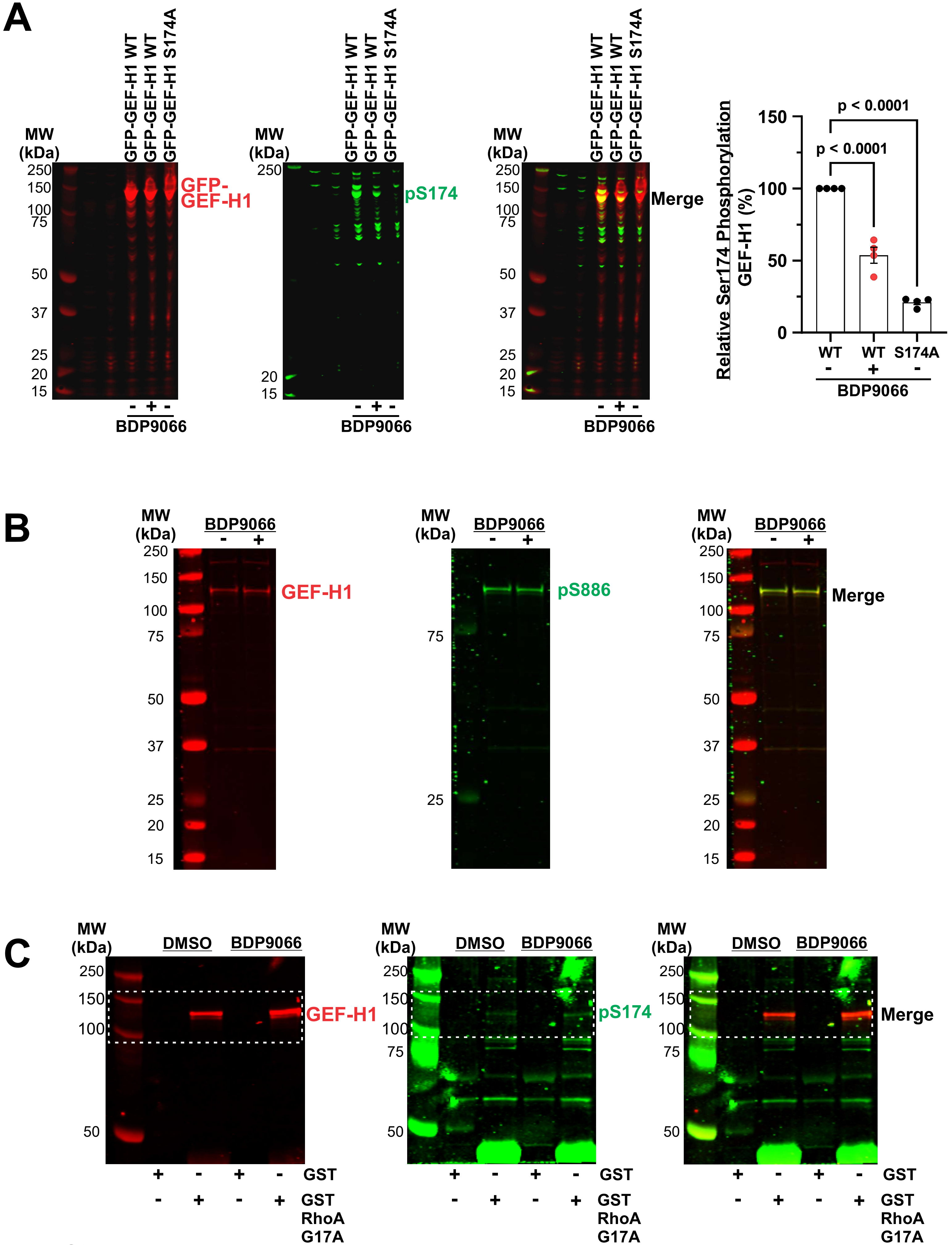

A
MW (kDa)
GFP-GEF-H1 WT
GFP-GEF-H1 WT
GFP-GEF-H1 S174A
250
150
100
75
50
37
25
20
15
GFP-GEF-H1
pS174
Merge
- + -
BDP9066
Relative Ser174 Phosphorylation GEF-H1 (%)
150
100
50
0
p < 0.0001
p < 0.0001
WT
WT
S174A
- + -
BDP9066
B
MW (kDa)
BDP9066
- +
GEF-H1
pS886
Merge
C
MW (kDa)
DMSO
BDP9066
GEF-H1
pS174
Merge
+ - + - GST
- + - + GST RhoA G17A


Figure S8

**Figure S8. MRCK phosphorylation of GEF-H1 on Serine 174. A.** *Left panel:* HEK293T cells were either non-transfected (NTC) or transfected with GFP-tagged wild-type (WT) or S174A mutant GEF-H1 that were treated with DMSO (-) or 10 µM BDP9066 (+) for 1 h, as indicated. Cell lysates were quantitatively western blotted with antibodies against GFP-epitope tagged GEF-H1 (red) and S174 phosphorylated GEF-H1 (pS174, green). *Right panel:* The ratios of pS174 to total GEF-H1 in each condition were normalized to DMSO cells expressing GFP-tagged GEF-H1 for each independent replicate. Means ± SEM, N = 4. Statistical analysis used one-way ANOVA with Tukey's multiple comparison test between indicated conditions producing the displayed *p* values. **B.** OVCAR8 cells were treated with DMSO vehicle control (-) or 10 µM BDP9066 (+) for 1 h, as indicated. Cell lysates were western blotted with antibodies against GEF-H1 (red) and Ser886 phosphorylated GEF-H1 (pS886, green). **C.** White rectangles with dotted borders indicate cropped regions used in Figure 7I.

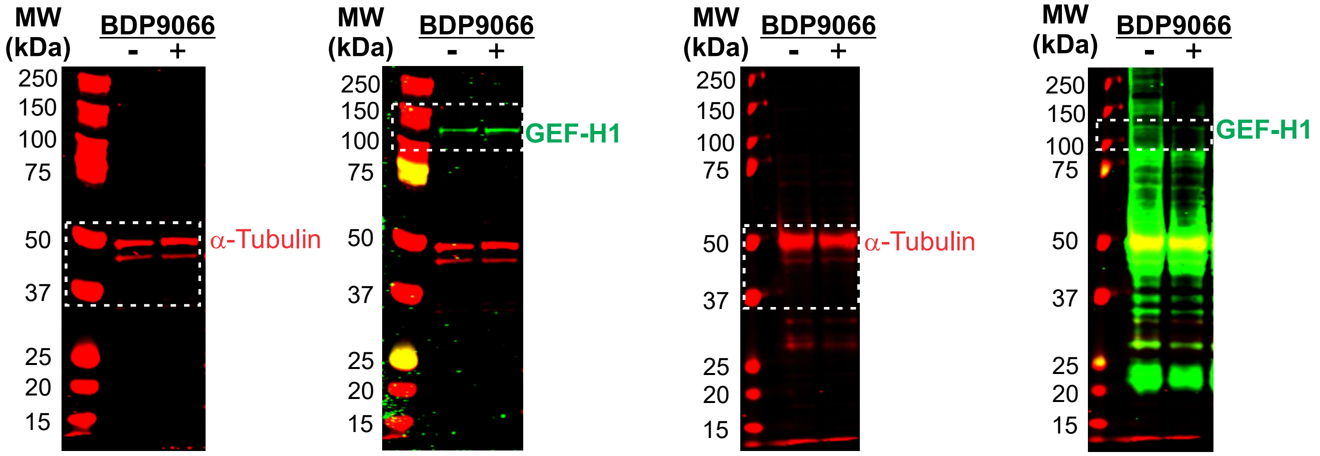

MW (kDa)
BDP9066
- +
250
150
100
75
50
37
25
20
15
α-Tubulin
MW (kDa)
BDP9066
- +
250
150
100
75
50
37
25
20
15
GEF-H1
MW (kDa)
BDP9066
- +
250
150
100
75
50
37
25
20
15
α-Tubulin
MW (kDa)
BDP9066
- +
250
150
100
75
50
37
25
20
15
GEF-H1


**Figure S9**

**Figure S9. MRCK inhibition reduces GEF-H1 association with α-Tubulin.** White rectangles with dotted borders indicate cropped regions used in Figure 8A.

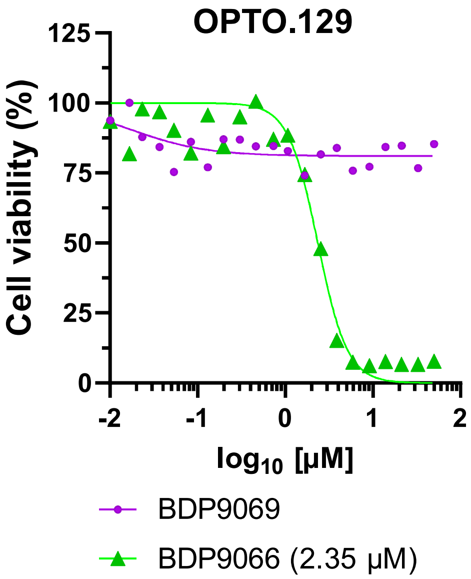

OPTO.129
Cell viability (%)
125
100
75
50
25
0
-2
-1
0
1
2
log10 [µM]
BDP9069
BDP9066 (2.35 µM)


**Figure S10**

**Figure S10. Decreased patient derived organoid viability induced by BDP9066 was not observed for the less potent MRCK inhibitor BDP9069 enantiomer.** The HGSOC patient derived organoid OPTO.129 grown on Matrigel was treated with DMSO vehicle control or indicated concentrations of active enantiomer BDP9066 or the less potent enantiomer BDP9069. Relative $EC_{50}$ values were calculated from 21-point drug concentrations of log(drug) versus response using four-parameter variable slope nonlinear curve fitting.

**Video 1. Time-lapse microscopy of non-targeting control (NTC) siRNA transfected OVCAR8 cells treated with BDP9066**. Related to Figures 9 A-B. A pair of NTC siRNA–transfected OVCAR8 cells was imaged for 14 min before addition of 3 μM BDP9066.

**Video 2. Time-lapse microscopy of GEF-H1-targted siRNA transfected OVCAR8 cells treated with BDP9066.** Related to Figures 9 A-B. GEFH1-depleted OVCAR8 cells were imaged for 14 min before 3 μM BDP9066 addition.

**Video 3. Time-lapse microscopy of NTC transfected OVCAR8 cells treated with DMSO.** Related to Figure 9B. A pair of NTC siRNA–transfected OVCAR8 cells was imaged for 14 min before addition of DMSO (0.1%).
